# Eyes on the prize: Mice deploy task-driven saccades during naturalistic foraging

**DOI:** 10.64898/2026.06.29.735201

**Authors:** Robert Taylor, Muad Y. Abd El Hay, Mina Glukhova, Cora Wolter, Marieke L Schölvinck, Martha N Havenith

## Abstract

Active sensing allows organisms to shape incoming sensory information through self-generated actions, and saccadic eye movements provide a key readout of this process in vision. In primates, saccades are strongly modulated by cognitive variables such as uncertainty, value, and behavioural goals, particularly in complex, naturalistic settings. In rodents, by contrast, saccades have largely been interpreted as reflexive or compensatory, and evidence that they are modulated by cognitive state, such as trial outcome, expectation, or task demands, has remained sparse. Here, we examined how mice use saccades during a vision-dependent foraging task in a traversable immersive virtual environment with naturalistic stimuli. We correlated their saccade dynamics with behavioural strategies observed within the virtual environment. Target-directed saccades emerged specifically when informative sensory evidence was available, but also occurred anticipatorily when animals could rely on previously learned spatial contingencies, indicating that saccades were guided not only by immediate visual input but also by internal representations. Strikingly, the temporal structure of inter-saccade intervals resembled signatures previously reported in primates and lengthened with increased processing demand following changes in task contingencies. Together, these findings show that mouse saccades are not merely reflexive gaze corrections, but form part of a cognitively modulated active sampling strategy. More broadly, they suggest that key principles of active visual sensing may be conserved across species and establish mouse oculomotor behaviour as a tractable readout of internal cognitive state.

## Introduction

Any interaction between an organism and its environment relies on a constant dynamic feedback loop linking sensation, perception, decision making and action (1). Vitally, this process generally involves active sensing, i.e. the active orientation of sense organs towards relevant stimuli, e.g via eye movements (2), whisking, sniffing, or touch (3). These active sensing behaviours do not simply provide a fixed or reflexive stimulus response; instead, they are modulated by ongoing cognitive processes such as search strategies (4), uncertainty (5) and attention (6).

In primates, active sensing is often studied in the form of saccadic eye movements, i.e. rapid gaze shifts used to sample the visual environment. The dynamics of saccades and gaze in macaques, including response times, sampling choices, and the allocation of fixations, are shaped by cognitive variables such as the subject’s internal model of environmental value (7), working memory (8), and uncertainty about the current decision (9). At the neural level, parietal cortex encodes the expected information gain of an impending saccade (5), indicating that the oculomotor system is dynamically tuned by cognitive state rather than operating as a fixed visuomotor reflex. Such internally generated dynamics emerge most clearly under naturalistic conditions, where the animal must flexibly coordinate its gaze with ongoing behavioural goals in a dynamic environment (10, 11). In more constrained paradigms such as simple fixation or cued saccade tasks, gaze can appear largely reflexive, obscuring the strategic, information-seeking character of gaze control that becomes apparent as sensory complexity and task demands approach those of the natural environment (12).

In contrast to the rich cognitive signatures of saccade dynamics observed in primates, the evidence for cognitive modulation of saccades in rodents is considerably more limited. While rodent studies have reported saccades comparable to those in primates in their basic kinematic properties (13, 14), the majority of eye movements observed in freely moving rodents appear to be tightly coupled to head movement, serving both to stabilize gaze during head tilt and, via a conjugate “saccade-and-fixate” pattern, to reset gaze in the direction of head rotation (15). A similar compensatory, saccade-and-fixate structure dominates eye movements even during active natural behaviours such as prey capture, where gaze shifts are largely tied to head movement rather than to the target (16). Even such gaze-resetting saccades appear to be driven by self-motion rather than directed toward specific visual targets. In head-fixed paradigms, where head-movement signals are absent, spontaneous saccades are infrequent and strongly influenced by motor state, being coordinated with navigational variables such as the direction of upcoming turns (17). To our knowledge there have been few reports of saccade dynamics modulated by cognitive state in rodents. Together, this body of evidence suggests that in rodents, saccades may mainly serve the largely automated (and at times predictive) compensation of visual drift in the presence of self-motion.

The apparent absence of significant cognitive modulation of mouse eye movements has mainly been met with two explanations. The first is that mice lack a foveal region of the retina, which has been interpreted as removing the benefit of reorienting gaze toward salient features. The second is that mice primarily rely on non-visual senses such as olfaction and whisking (18), rendering active visual sampling secondary or redundant.

Despite the prevailing view that, especially in head-fixed paradigms, mouse saccades are rare and primarily compensatory, the mouse visual system is in fact equipped for directed gaze. Topographic analyses of the mouse superior colliculus (SC) reveal a motor map for directed eye movements (19), mirroring the organisation of the primate oculomotor SC, and mouse V1 contains a region of improved visual processing analogous to a fovea (20), directly challenging the assumption that, lacking a retinal fovea, mice have little to gain from reorienting their gaze. This machinery is actively engaged: the mouse SC drives stimulus-evoked, directionally biased gaze shifts in head-fixed animals, with a flexibility that exceeds what a purely compensatory account would predict (21), and in free-viewing the typical amplitude of mouse saccades corresponds to the minimum gaze displacement required to decorrelate V1 population responses across successive fixations in natural scenes, a distance that scales with receptive field size (14). This principle generalises across mammals not as a fixed behaviour but as a common functional outcome: despite large differences in receptive-field size, different species achieve comparable neural coverage of the visual scene through distinct, species-specific sampling behaviours, consistent with eye movements being calibrated to the spatial grain of cortical representation (22). Consistent with an active sampling function, gaze shifts in freely moving mice initiate structured sequences of visual cortical processing, indicating that mouse saccades actively organise visual encoding rather than merely accompanying movement (23). Mice thus appear to position their eyes to maximise novel sensory input per fixation, rather than merely redirecting gaze to compensate for body movement, and they successfully rely on visual cues in naturalistic contexts such as evading predators (24, 25) and prey capture (26).

This evidence suggests that mouse vision is a more complex and actively deployed system than often assumed, but one whose dynamics may only become apparent when a task specifically demands active visual engagement. This implies that rich saccadic behaviour and its cognitive modulation in rodents may only manifest when the sensory environment and behavioural context supports and requires it. It is therefore possible that the reason state-dependent saccade dynamics documented in primates may not yet have been reported in mice is methodological rather than biological. Freely moving studies, while providing naturalistic conditions, have focused primarily on compensatory gaze stabilisation (15), and although head-fixed work has shown that mouse saccades can be stimulus-evoked and flexibly directed (21) and coordinated with navigational variables such as upcoming turns (17), and naturalistic freely-moving studies have characterised gaze during behaviours such as prey capture (16), these paradigms have rarely included a behavioural readout linking saccade dynamics to trial-by-trial task performance, outcome, or internal state.

Here, we test how mice employ saccades during a dynamic foraging task that requires vision, using free navigation within an immersive virtual environment featuring naturalistic stimuli. We find that mouse saccades are cognitively modulated; shaped by trial outcome, working memory, and task complexity in ways that closely parallel primate gaze behaviour. This provides, to our knowledge, the first direct investigation of mouse saccadic dynamics within an environment explicitly designed to probe active visual sensing under naturalistic conditions.

## Results

### A. Mice saccade during a virtual navigation task

We developed a virtual reality (VR) paradigm in which head-fixed mice run on an omnidirectional trackball to approach naturalistic visual stimuli projected onto a dome-shaped enclosure covering 250° of visual angle (Fig. 1A) (27–29). This approach enables a wide range of non-discrete behavioural responses, which are tracked via extensive behavioural read-outs from running trajectories to facial and posture markers, pupillometry and eye movement tracking to lick behaviour. Furthermore, the immersive nature of the visual environment, featuring naturalistic stimuli and self-initiated visual flow, enables rapid task learning (<5 sessions to train visual discrimination, (27)).

**Fig. 1.**
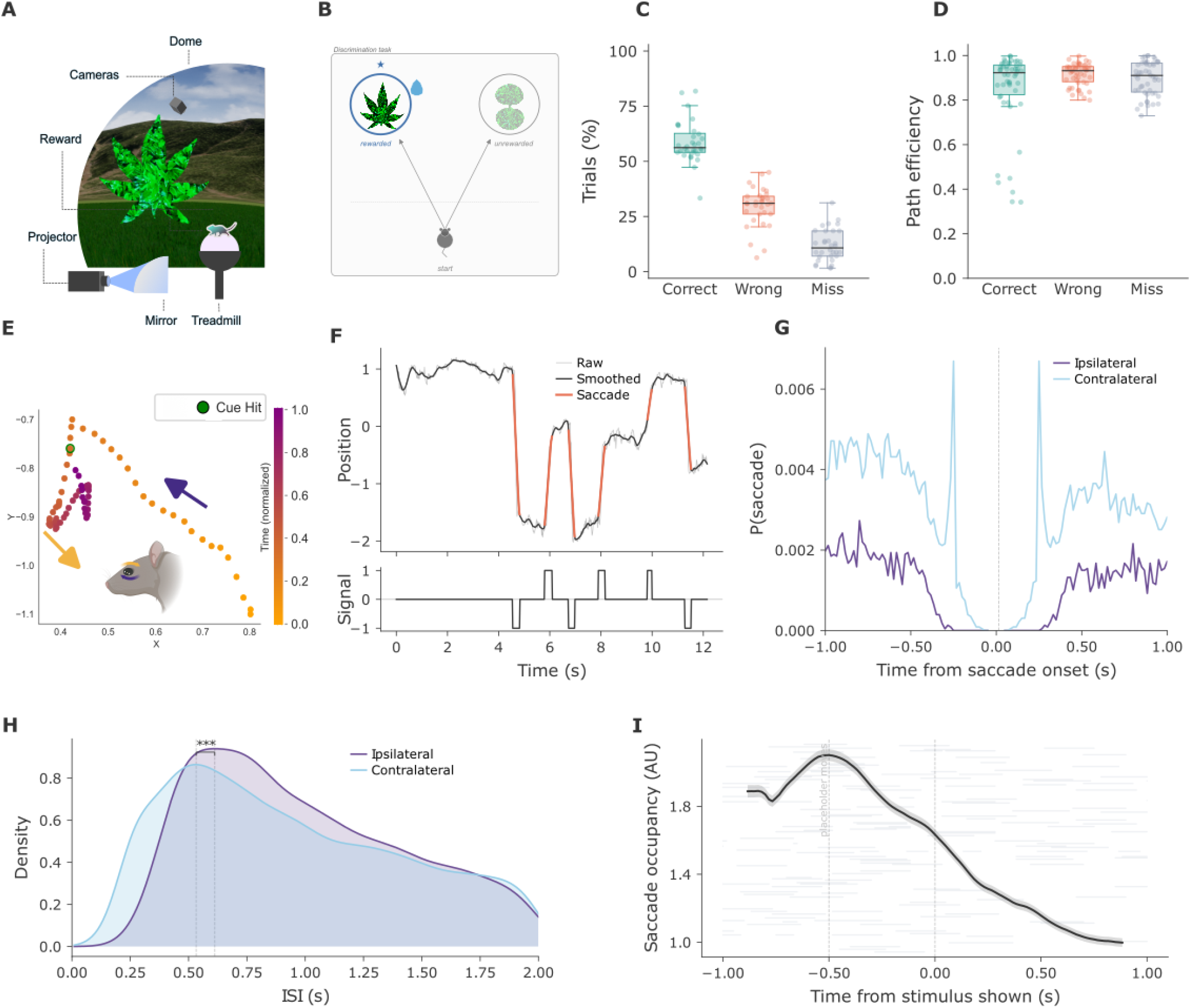
Behavioural task design and eye movement characterisation in head-fixed mice. **A)** Schematic of the experimental apparatus. **B)** Task structure. An ambiguous placeholder appeared centrally; upon approach it split into a laterally offset stimulus and distractor. Navigating to the stimulus yielded a reward; hitting the distractor triggered a timeout corridor. Misses restarted the trial. **C)** Trial outcome distributions across animals (n = 11). Box plots show proportions of correct (green), wrong (red), and miss (grey) trials. **D)** Path efficiency (optimal / actual path length) by trial outcome for pre-switch trials (box plots with median and 25th/75th percentiles, scatter displays individual sessions). **E)** Example LightningPose-labelled pupil trajectory in camera space. Coloured dots show time-normalised eye positions (0–1); arrows indicate detected saccades (purple = rightward, orange = leftward). **F)** Representative pupil position trace (top) and saccade detection signal (bottom). Raw (grey) and smoothed (blue) traces shown; detected saccades highlighted in orange. **G)** Peri-saccade probability aligned to saccade onset for ipsilateral and contralateral saccades. Note the stereotyped contralateral return to baseline at ^*∼*^250 ms.**H)** Inter-saccade interval peak density plot for ipsidirectional (dark purple, N = 4,646) and contradirectional (light blue, N = 20,178) saccades (log-density axis). **I)** Peri-stimulus saccade raster and PSTH aligned to stimulus onset (N = 14,142 trials). Grey: individual saccade events, each row depicts one trial; black: population moving sum (window = 15 frames). Dashed line = stimulus onset.

On each trial, mice initiated navigation toward a centrally presented ambiguous ‘placeholder’ stimulus, which then split into a rewarded target stimulus and an unrewarded distractor stimulus, both placed at a lateral offset (Fig. 1B). All stimuli were modelled on naturalistic leaf shapes. Animals were trained to approach the target stimulus in order to obtain food rewards in the form of sweetened oat milk (correct trials); distractor contact triggered a timeout in a corridor the mice had to traverse before restarting the same trial (wrong trials); and failure to make contact with either target or distractor (miss trials) caused the trial to restart without a preceding timeout corridor.

Across the cohort (n = 11 animals), group-level accuracy was 57.9% ± 2.4% correct, with wrong responses accounting for 29.7% ± 1.6% and misses for 12.4% ± 1.8% of trials (mean ± SEM; Fig. 1C). Note that due to the open spatial layout of the task, which does not force stimulus choices and requires animals to actively navigate away from their central running trajectory in order to approach a stimulus, random success rate was extraordinarily low. In a closely related version of the same virtual-environment task, removing visual input abolished directed responses entirely, with animals ceasing to change running direction in the absence of a visible target, resulting in a chance success rate of zero (28). As such, target approaches in this paradigm reflect deliberate, visually guided choices rather than random navigation, and the observed accuracy lies significantly above any uninformed baseline.

In addition to basic trial outcomes, we also quantified two further markers of effective task performance: (1) Reaction times were determined based on the point of strongest inflection in running direction, marking the time at which animals began to actively navigate to their chosen stimulus (Fig. S1, see Methods). (2) Path efficiency was computed as the ratio between the actual length of an animal’s running trajectory post stimulus onset, and the theoretical shortest path towards its chosen stimulus. As such, it reflects the presence or absence of additional direction changes or detours in the animal’s target approach.

Descriptively, path efficiency tended to be lower and reaction times longer in correct than in wrong or miss trials (Fig. 1D, S1), consistent with animals exhibiting more deliberate, exploratory behaviour as they searched for correct stimuli, while behaviour on wrong and miss trials appeared more stereotyped and direct.

To explore if these varying approaches to task performance were also reflected in active sensing strategies, we tracked eye movements throughout the task. To this end, we used LightningPose (30) to extract pupil position from bilateral video recordings of each eye, acquired at a frame rate of 60 Hz (Basler acA640-121gm; see Methods). A single LightningPose model was trained across animals and sessions to track both eyes (Fig. 1E). To select sessions with reliable pupil estimates, we used the per-frame, per-keypoint confidence scores returned by LightningPose (range 0–1), averaged across all frames and keypoints within each session. Sessions with a mean confidence score above 0.97 were retained. This criterion yielded 140 of 205 sessions from 11 animals (median 11 sessions per animal, range 3–27). In the selected data sets, saccades were detected from the smoothed pupil position signal using a velocity thresholding algorithm (Fig. 1F, see Methods).

In the selected data sets, we first established to what extent saccades occurred simultaneously or independently across both eyes. Head-fixed mice displayed highly synchronised eye movements between eyes (Fig. S2). Based on this result, we restricted our subsequent analyses to videos of the left eye, since tracking quality was marginally higher (see Methods), and eye movements extracted from one eye were set to faithfully capture binocular saccade dynamics.

Next, we examined the temporal structure of saccade sequences. As the cross-correlation histogram of saccades shown in (Fig. 1G) demonstrates, there was a marked tendency for saccades to be followed by a contralateral saccade at a rather specific delay of approx. 250 ms, suggesting that at least some saccades follow a temporally stereotyped pattern. This pattern did not occur for ipsilateral saccades, i.e. saccades moving in the same direction as the previous one. The peak in the occurrence of contralateral saccades at a delay of *∼*250 ms is broadly consistent with the *∼*200 ms minimum intersaccadic interval reported in humans (31) and falls within the 2–6 Hz saccade-rate range (mean intersaccadic intervals 240–270 ms) reported for non-human primates (32). Consistent with this observation, inter-saccade intervals were significantly shorter for contradirectional than ipsidirectional saccades (Fig. 1H; Mann–Whitney *U* = 5.08 × 10^7^, *n* = 4,646 ipsidirectional / 20,178 contradirectional saccades, median 0.900 vs. 0.850 s, *p* = 2.9 × 10^−19^, CLES = 0.542, rank-biserial *r* = +0.08).

Finally, we characterised the timing of saccades relative to the time course of the task. Aligning saccade events to stimulus onset revealed that saccades occurred most frequently around the appearance of the initial ‘placeholder’ stimulus, i.e. shortly before the appearance of target and distractor. Saccade rate then continued to decline once animals were moving towards their chosen stimulus (Fig. 1I, Fig. S6). Together, these results establish that during virtual navigation, saccadic eye movements in mice follow a clear temporal structure and are modulated by task events. This in turn poses the question what functions such task-modulated saccades might serve.

### B. Saccades are not closely tied to motor preparation

A prevailing view holds that saccades in rodents largely serve gaze stabilisation during head movements (15), and, in head-fixed settings, are coordinated with navigational motor planning such as the direction of upcoming turns (17). Building on this, we first asked whether saccades in our paradigm were tied to locomotor activity, for instance occurring around movement onset. Running speed showed only a modest increase after saccades (Fig. 2A). Although post-saccade running speed differed significantly from a shuffled baseline (Fig. 2B; Mann–Whitney *U* = 1.94 × 10^9^, *n* = 61,614 / 61,553, median 7.96 vs. 8.01 cm/s, *p* = 3.1 × 10^−10^, CLES = 0.510, rank-biserial *r* = +0.02), the magnitude of this difference was negligible and saccades occurred across the entire range of running speeds, arguing against a tight, motor-preparatory coupling between saccades and movement onset. Having established that saccades are not consistently time-locked to locomotor activity, we next examined whether they were instead coupled to discrete navigational events, specifically to changes in running direction, as reported previously for head-fixed mice (17). To test this relation, we computed the latency between each saccade and the largest turn (i.e. change in running direction) per trial. While these maximal direction changes did preferentially occur around saccade onsets (Fig. 2C), saccades and direction changes were observed at a wide variety of delays, with saccades leading direction changes in some cases, and lagging in others. This suggests that while saccades do indeed often co-occur with heading changes, their temporal relationship is highly varied, arguing against a reflexive or otherwise stereotyped link between them.

**Fig. 2.**
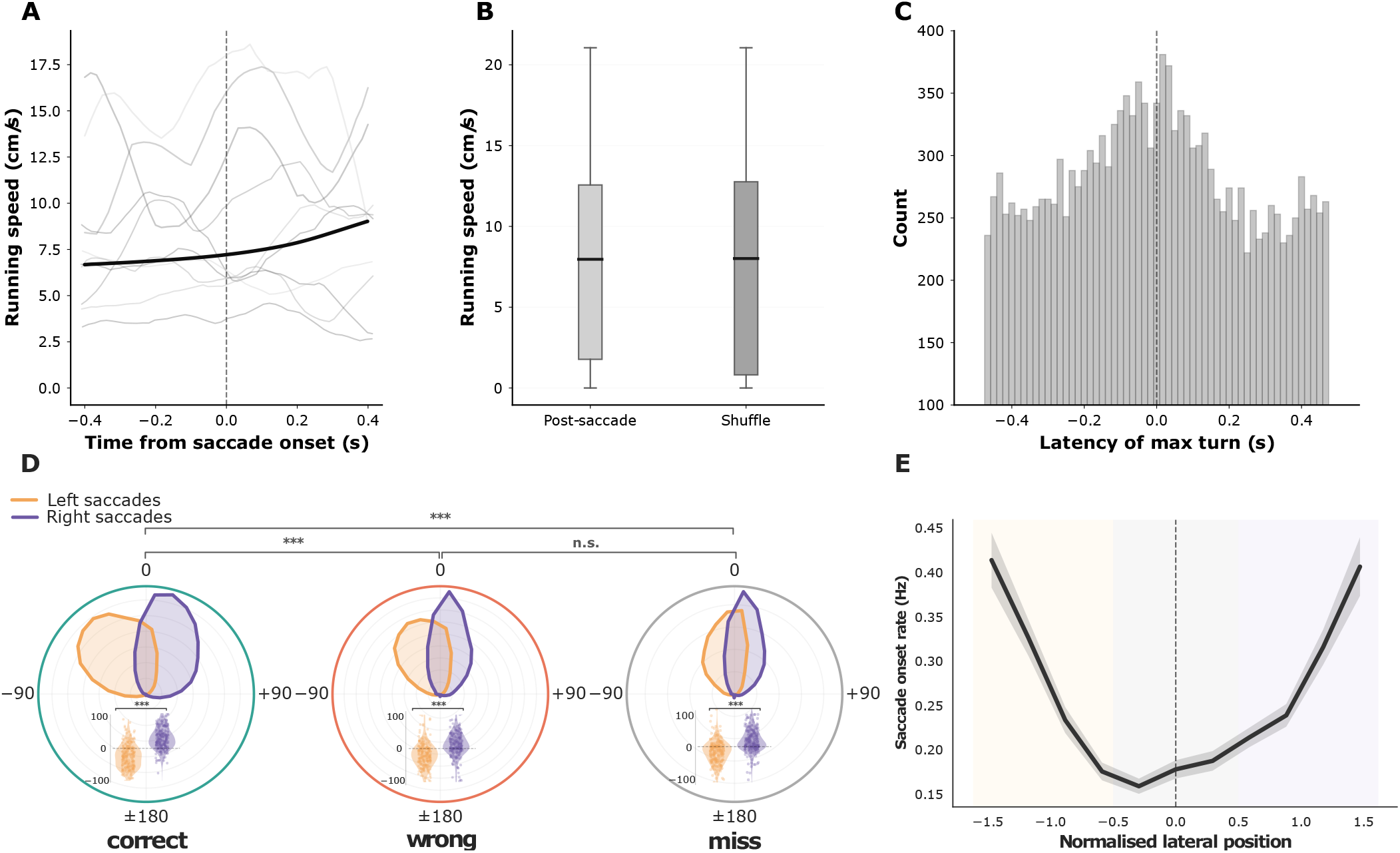
Saccade-related locomotor dynamics and spatial dependencies in head-fixed mice behaving in a virtual reality task. **A)** Saccade-triggered average running speed (cm/s, mean ± s.d.) aligned to saccade onset (dashed line; N = 62,845 saccades, 11 animals, 140 sessions). Grey lines: 5 example traces **B)** Running speed (cm/s) in a 0–0.5 s post-saccade window versus shuffled non-saccade timepoints. Box plots show median, IQR, and 1.5× IQR (N = 61,553). **C)** Latency from saccade onset to maximal turn amplitude. Dashed line = saccade onset. **D)** Post-stimulus turn-change angle distributions by saccade direction and outcome. Inset: signed turn-change amplitude by saccade direction and outcome.**E)** Saccade onset rate (Hz) by normalised lateral position (N = 25,514; shading = SEM). Inset: mean ± SEM for Left, Centre, and Right regions; dots = sessions.

This notion was also supported when comparing not the timing but the targeted direction of saccades and locomotor activity. Fig. 2D shows the change in running direction around each saccade, computed as the difference between mean heading over the 0.5 s preceding saccade onset and the 1.0 s following it, for rightward (purple) and leftward (orange) saccades, presented separately for correct, wrong and miss trials. These distributions indicate that running directions tend to shift towards the direction of preceding saccades, suggesting a somewhat preparatory role for saccades. This directional coupling was present across all trial outcomes but was strongest on correct trials, where running directions following leftward versus rightward saccades showed limited overlap. This coupling between saccade and turn direction was significant on correct, wrong, and miss trials (Fig. 2D; Mann–Whitney tests of rightward vs. leftward saccades on signed turn: correct *p*< 10^−300^, CLES = 0.857, *r* = +0.71; wrong *p* = 1.6 × 10^−66^, CLES = 0.731, *r* = +0.46; miss *p* = 1.8 × 10^−58^, CLES = 0.738, *r* = +0.48; BH-corrected). Cross-outcome comparison of turn magnitude showed larger turns on correct than wrong or miss trials (correct vs. wrong *p* = 1.4 × 10^−25^, CLES = 0.585, *r* = +0.17; correct vs. miss *p* = 2.7 × 10^−30^, CLES = 0.600, *r* = +0.20; wrong vs. miss n.s., *p* = 0.19, *r* = +0.03). These results emphasise that while saccades are often directed at parts of the visual scene that animals are about to approach, this relation is not set in stone, and can be modulated by context.

Finally, while the relation between saccade and running direction was modulated by trial outcome, it was not significantly impacted by saccade amplitude, with leftward and rightward saccades consistently associated with leftward and rightward turns irrespective of saccade size (Fig. S3). This result further argues against a tight, amplitude-graded mechanical coupling between saccades and changes in running direction.

Next, to test whether saccades in this paradigm could be interpreted as information-seeking behaviour, we examined where in the virtual task environment they were initiated. To this end, we mapped saccade likelihood depending on the animal’s lateral position relative to their chosen stimulus. As shown in (Fig. 2E), the occupancy-normalised saccade onset rate was highest when animals were positioned laterally from their chosen stimulus, and lowest when they were directly in front of it, with a pronounced suppression at central positions and a symmetric rise toward either flank. This pattern also held when mapping preferred saccade onset locations across the entire virtual task space, with saccades concentrated laterally from stimulus locations across the entire length of the task space, with relative suppression along the central corridor leading directly toward the stimulus (Fig. S4).

Finally, the distribution of saccade locations was also modulated by trial outcome: In correct trials, saccades were predominantly directed toward the chosen stimulus across a broad range of locations, whereas in wrong trials, a substantial fraction of saccades were instead directed away from the chosen stimulus, particularly at later approach positions (Fig. S5). These results indicate that saccades are deployed selectively when behavioural changes are needed to achieve targeted outcomes, and that this mechanism operates more effectively in correct than wrong trials. This suggests that saccades are actively deployed during locomotion, based on the animal’s position relative to task-relevant stimuli, necessary heading corrections as well as predicted trial outcomes, a pattern that is consistent with a role for saccades in actively guiding behaviourally relevant navigational decisions.

### C. Saccade structure predicts trial outcome

Based on the observation that both saccade location and coupling to locomotion are modulated by trial outcome, we next asked whether saccade directions could be directly predictive of the animal’s upcoming stimulus choices. Overall saccade rate decreased gradually as the trial progressed for both correct and wrong trials (Fig. 3A). This tallies with saccadic inhibition observed in freely viewing macaques, in which the onset of a visual stimulus transiently suppresses saccades (33). Saccade rate did not differ significantly between correct and wrong trials at the session level, either before or after stimulus onset (Fig. 3B; session-level Wilcoxon signed-rank, prestimulus *p* = 0.052, post-stimulus *p* = 0.062, *n* = 32 sessions; both n.s. after correction), though wrong trials showed a numerical tendency toward higher rates.

**Fig. 3.**
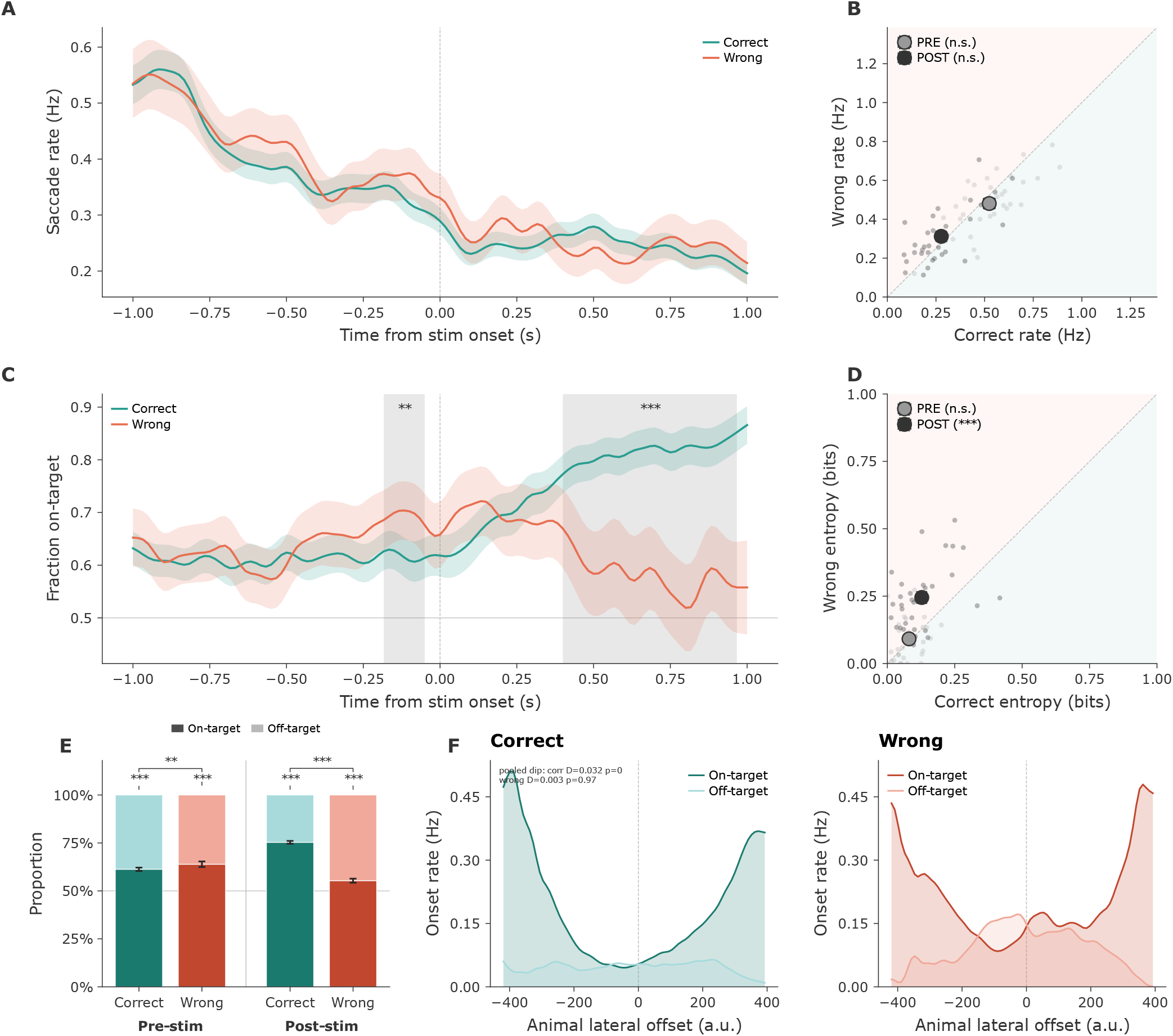
Saccade orienting behaviour predicts trial outcome in a freely moving visual discrimination task. Statistical details are given in the Results; saccade rate and entropy comparisons are evaluated at the session level, all other comparisons on pooled saccades. **A)** Saccade rate over time (correct, green; wrong, red; 31,454 correct / 10,866 wrong trials, 11 animals, 140 sessions). Shading = binomial standard error (pooled). **B)** Pre-stimulus (grey) and post-stimulus (black) saccade rate, correct versus wrong trials (32 sessions, 8 animals). **C)** Fraction of saccades directed toward the chosen stimulus over time (correct, green; wrong, red; 31,454 correct / 10,866 wrong trials). Shading = binomial standard error (pooled); grey bar marks the significant cluster (see Results). **D)** Pre-stimulus (grey) and post-stimulus (black) saccade entropy, correct versus wrong trials (32 sessions, 8 animals). **E)** Proportion of pre- and post-stimulus saccades toward versus away from the chosen stimulus (pre-stimulus *n* = 11,861 correct / 4,156 wrong; post-stimulus *n* = 13,136 correct / 8,593 wrong saccades). Dashed line = chance. **F)** Occupancy-normalised saccade onset rate as a function of the animal’s lateral position relative to the chosen stimulus, shown separately for on-target and off-target saccades on correct (left) and wrong (right) trials. Rate is onsets per unit dwell time per position bin (see Methods).

In contrast, the directional structure of saccades diverged noticeably. In correct trials, the fraction of saccades directed toward the stimulus the animal would ultimately choose (termed ‘on-target saccades’) rose sharply after stimulus onset, whereas in wrong trials, it remained close to chance (Fig. 3C; cluster-based permutation test, significant post-stimulus cluster spanning 0.400–0.967 s, Σ*z* = 225.1, *p*< 0.0002). A smaller cluster shortly before stimulus onset (− 0.183 to −0.050 s, Σ*z* = −23.8, *p* = 0.0024) reflected a brief difference in the opposite direction, with a marginally higher on-target fraction on wrong than correct trials (69.9% vs. 61.6%) just before the stimulus divided. Across all post-stimulus saccades, 75.3% of saccades in correct trials were on-target, compared with 55.3% in wrong trials. The post-stimulus on-target fraction exceeded chance on both correct and wrong trials, and was substantially higher on correct trials (Fig. 3E; binomial test vs. 0.5: correct 9,897/13,136 = 0.753, Cohen’s *g* = +0.253, *q*< 10^−300^; wrong 4,756/8,593 = 0.554, *g* = +0.053, *q* = 4.4 × 10^−23^; correct-versus-wrong *χ*^2^ *q* = 5.2 × 10^−207^, Cramér’s *V* = 0.209, odds ratio 2.47). This indicates that saccadic orienting toward the chosen stimulus emerges rapidly once relevant visual information is present, and is a robust predictor of successful outcomes.

This conclusion also applies to the overall temporal structure of saccades across trial types. To capture this temporal structure, we computed saccade entropy, which is a metric reflecting the random dispersion of saccade directions across the time course of the trial. Saccade entropy was significantly lower in correct than wrong trials in the post-stimulus period (Fig. 3D; session-level Wilcoxon signed-rank, *p* = 5.0 × 10^−5^, *n* = 32 sessions, *r* = +0.82; higher on wrong trials in 26 of 32 sessions). This suggests that correct trials are characterised by more predictable, directed eye movements, whereas wrong trials contain more random visual scanning. Finally, we examined how the rate of on-target and off-target saccades depended on the animal’s lateral position relative to the chosen stimulus. To control for the uneven time animals spent at each lateral position, saccade onsets were expressed as an occupancy-normalised rate (onsets per unit dwell time in each position bin; see Methods). On correct trials, on-target saccades were strongly suppressed when the animal was centred on the chosen stimulus and rose sharply as it became laterally offset to either side, producing a pronounced bimodal, U-shaped dependence on position (Hartigan’s dip test on the occupancy-normalised distribution: correct *D* = 0.032, *p*< 0.001, *q* = 0.002, bimodal; wrong *D* = 0.003, *p* = 0.97, *q* = 0.97, unimodal; Fig. 3F); off-target saccades, by contrast, occurred at a low, roughly uniform rate across positions. On wrong trials this position dependence was far weaker, and off-target saccades became more prominent at central positions. In other words, on correct trials saccades were recruited specifically when the animal was spatially offset from its target and were predominantly directed toward it, whereas on wrong trials saccades were deployed less selectively with respect to position and more often directed away from the chosen stimulus.

Together, these results demonstrate that saccade behaviour is not merely correlated with trial outcome but reflects a qualitatively different gaze strategy: correct trials are characterised by directed, spatially structured orienting toward the target, whereas wrong trials show unpredictable scanning with weaker directional bias and lower navigational coupling.

### D. Saccade orienting reflects working memory

While the saccades analysed so far show clear modulation by task context, they always occur in the presence of visual target stimuli, and can therefore be seen as a cognitively modulated reaction to these stimuli. To test if saccades could also be deployed in a task-relevant way in the absence of relevant visual stimuli, we tested if animals showed anticipatory gaze strategies in trials where the location of the target stimulus was known beforehand.

For training purposes, our task included a post-error condition, whereby after an incorrect response, the same trial was repeated until the animal responded correctly (Fig. 4A; see Methods). Such corrective post-error trials thus allowed animals to predict the rewarded stimulus location based on the previous trial outcome.

**Fig. 4.**
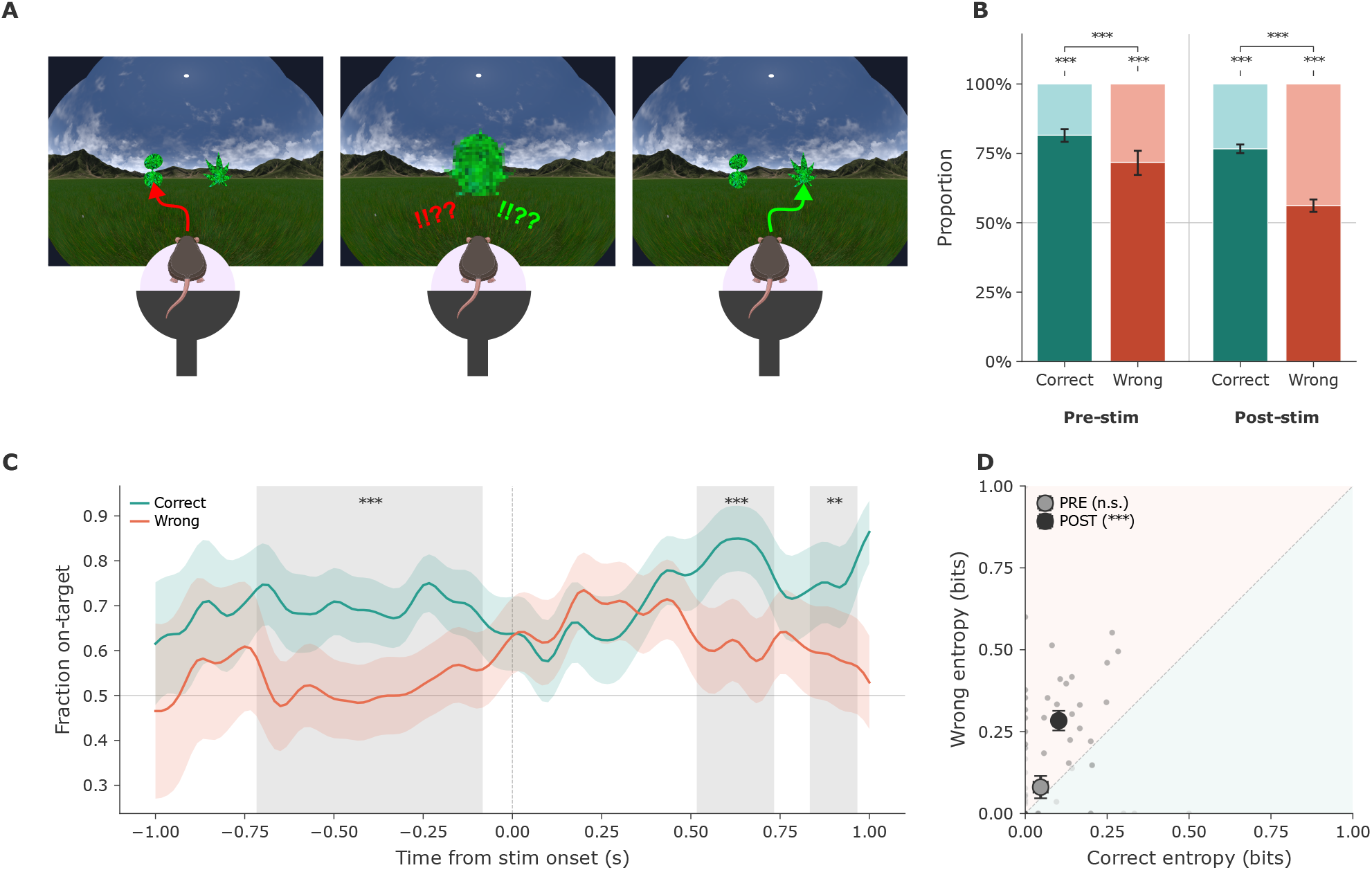
Saccade orienting behaviour predicts spatial contingencies. Statistical details are given in the Results. **A)** Schematic describing post-error trials (see Methods). **B)** Proportion of pre- and post-stimulus saccades toward versus away from the chosen stimulus, for post-error trials only (pre-stimulus *n* = 1,116 correct / 411 wrong; post-stimulus *n* = 2,816 correct / 1,875 wrong saccades). Dashed line = chance. **C)** As in Fig. 3C, for post-error trials only. The correct-versus-wrong divergence in on-target fraction was significant both before and after stimulus onset (see Results); shading = SEM. **D)** As in Fig. 3D, for post-error trials only (entropy, session level).

Here, we computed the fraction of on-target saccades around stimulus onset for post-error trials (Fig. 4B). On correct post-error trials, the on-target fraction was significantly above chance even before stimulus onset (Fig. 4B; prestimulus, 910/1,116 = 0.815 on-target, binomial test vs. 0.5, Cohen’s *g* = +0.315, *q* = 1.7 × 10^−105^), and exceeded the corresponding wrong-trial fraction (pre-stimulus wrong 295/411 = 0.718, *g* = +0.218; correct-versus-wrong *χ*^2^ *q* = 4.5 × 10^−5^, Cramér’s *V* = 0.106). This divergence was already significant before stimulus onset and was maintained after it (post-stimulus correct 0.767, wrong 0.562; *χ*^2^ *q* = 4.1 × 10^−49^, Cramér’s *V* = 0.216, odds ratio 2.57), with the correct-versus-wrong divergence reaching significance both before and after stimulus onset (Fig. 4C; cluster-based permutation, pre-stimulus cluster −0.717 to −0.083 s, Σ*z* = 137.6, *p*< 0.0002; post-stimulus clusters 0.517–0.733 s, Σ*z* = 51.8, *p*< 0.0002, and 0.833–0.967 s, Σ*z* = 24.0, *p* = 0.0016). That the on-target bias is present before the discriminative stimulus appears, on trials where the rewarded location can be inferred from the previous trial, indicates that gaze is directed in anticipation of a known spatial contingency rather than purely in reaction to the visual scene. By contrast, on regular trials, where the rewarded location cannot be known until the stimulus divides, the target-directed (correct > wrong) divergence emerged only after stimulus onset (Fig. 3C): the small pre-stimulus cluster on regular trials ran in the opposite direction (wrong > correct): before the stimulus divided, gaze on error trials was already biased toward the stimulus the animal would go on to choose, rather than reflecting anticipatory orienting toward the rewarded location. The anticipatory, target-directed pre-stimulus bias was thus specific to post-error trials, confirming that it arises only when the rewarded location is predictable.

While correct post-error trials showed an anticipatory directional bias toward the target before stimulus onset (Fig. 4C), the predictability of gaze (entropy) did not yet differ between correct and wrong trials at this stage (session-level Wilcoxon, pre-stimulus *p* = 0.25, *n* = 30 sessions). Only once stimuli appeared did saccadic entropy rise on wrong trials relative to correct (Fig. 4D; session-level Wilcoxon signed-rank, post-stimulus *p* = 2.3 × 10^−5^, *n* = 31 sessions, *r* = +0.87; higher on wrong trials in 26 of 31 sessions).

Taken together, these results dissociate two components of saccadic behaviour and reveal a predictive gaze strategy in mice: a pre-stimulus bias toward the anticipated target, reflecting spatial contingencies held in memory before stimulus onset, and a post-stimulus increase in saccadic disorganisation on wrong trials that emerges only once visual stimuli are present. This anticipatory, memory-driven process is thus dissociable from the stimulus-evoked scanning behaviour observed during active visual search.

### E. Saccade strategy becomes more variable after rule switching

To further explore how task demands and stimulus features drive saccades, we next tested how saccades might shift when the same stimuli were associated with new behavioural outcomes. To this end, we introduced uncued switches in stimulus contingencies: an intermediate stimulus was introduced, which could either serve as the target or distractor depending on which stimulus it was compared against (Fig. 5A; see Methods). The switch between stimulus contingencies always happened after 10 correctly responded trials (5–20 trials in total, median 12, depending on task performance). This design allowed us to test whether more structured saccadic behaviour predicted successful task performance also as task rules changed, or whether saccade patterns reflected previously learned reward contingencies. Overall task performance was comparable before and after rule switches (pre-switch: 17 sessions; post-switch: 90 sessions; Fig. 5B). The per-animal proportion of correct trials did not differ significantly across the switch (Wilcoxon signed-rank, *n* = 9 animals, *W* = 8, *p* = 0.098, rank-biserial *r* = 0.64; correct 63.3% → 69.2%), and neither wrong-nor miss-trial rates changed significantly (*p* = 0.82 and *p* = 0.16, respectively). In other words, animals switched between task rules without a significant disruption to overall accuracy. However, this does not imply that behaviour was entirely unaffected by rule switches. Path efficiency decreased significantly across the switch for both outcomes (Fig. 5C; Mann–Whitney, pre-vs. post-switch: correct median 0.958 → 0.922, *p* = 2.5 × 10^−11^, *r* = +0.11; wrong median 0.948 →0.898, *p* = 4.9 × 10^−29^, *r* = +0.29), with the larger shift on wrong trials. Approaches thus became less direct after the switch, particularly on error trials.

**Fig. 5.**
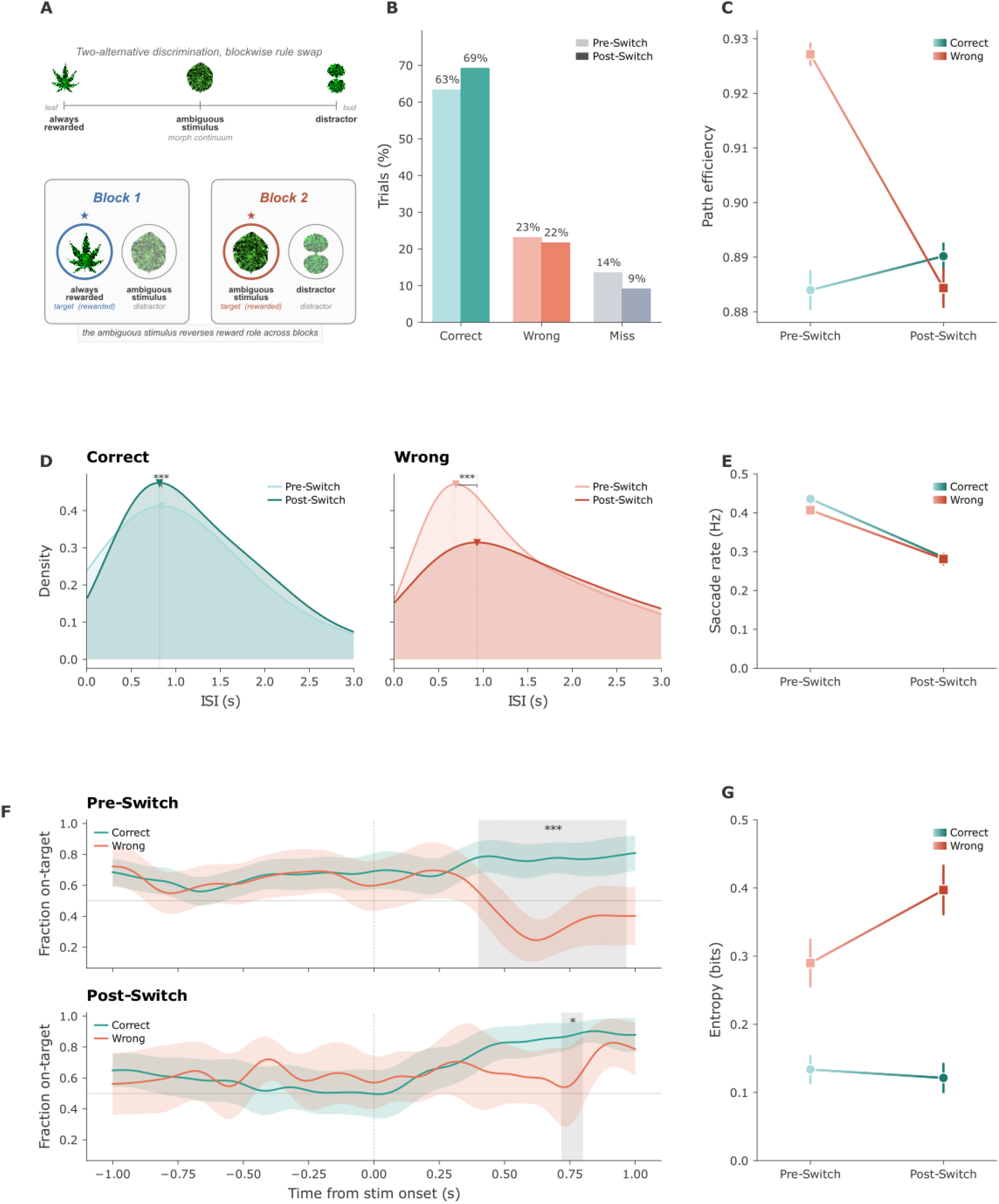
Saccade orienting strategy is preserved but becomes more variable after a stimulus identity switch. Statistical details are given in the Results; performance (B) is evaluated at the animal level and entropy (G) at the session level, all other comparisons on pooled trials. **A)** Schematic of the stimulus switch manipulation. Animals learned a two-choice discrimination, then generalised across morphs until a 50% morph served as either S+ or S−. In the switch block, the role of the 50% morph alternated in blocks of correct trials (typically 10). **B)** Performance before (pre-switch) and after (post-switch) the switch (*n* = 9 animals). Stacked bars show correct (green), wrong (red), and miss (grey) proportions (pre-switch: 17 sessions, 4,772 trials; post-switch: 90 sessions, 1,737 trials). Pre-switch trials comprise the full pre-switch sessions, whereas post-switch performance is evaluated on the switch trials themselves; this performance trial set is distinct from the peri-switch windows used for the saccade analyses in C–G (see Methods). **C)** Path efficiency by outcome, pre-switch versus post-switch; arrows indicate direction of change. **D)** Inter-saccade interval distributions pre- and post-switch, shown separately for correct (left) and wrong (right) trials. Kernel density estimates (Silverman bandwidth); triangles mark mode. **E)** Post-stimulus saccade rate (Hz) by outcome, pre-versus post-switch. **F)** Fraction of on-target saccades over time for pre-switch (top) and post-switch (bottom); shading = SEM, grey bar marks the significant cluster. **G)** Post-stimulus saccade entropy by outcome, pre-versus post-switch.

If the rate of saccadic sampling in mice reflects the processing demands of the visual environment, as reported in primates (32), then the increased processing demand following a rule switch should be accompanied by a lower saccade rate and correspondingly longer inter-saccadic intervals. Inter-saccadic intervals lengthened significantly after the switch on both correct and wrong trials (Fig. 5D; Mann–Whitney, pre-vs. post-switch: correct median 0.933 →1.117 s, *p* = 1.7 × 10^−7^, *r* = −0.12; wrong median 1.175 →1.533 s, *p* = 1.4 × 10^−8^, *r* = −0.16; on full-trial ISIs). This was accompanied by a significant reduction in post-stimulus saccade rate across outcomes (Fig. 5E; Mann–Whitney, correct *p* = 9.7 × 10^−8^, *r* = +0.06; wrong *p* = 3.7 × 10^−7^, *r* = +0.10; display rates 0.44/0.41 → 0.29/0.28 Hz). The post-switch lengthening of fixation durations was therefore consistent across correct and wrong trials, in keeping with a global adjustment of saccadic timing to the increased processing demand of the post-switch environment.

We next examined whether the temporal dynamics of saccades were affected by rule switching. Before the switch, the fraction of on-target saccades diverged strongly between correct and wrong trials after stimulus onset (Fig. 5F, top; cluster-based permutation, 0.400–0.967 s, Σ*z* = 130.5, *p*< 0.0002, Fig. S7). After rule switching, this divergence was markedly reduced, with only a single brief significant cluster remaining (Fig. 5F, bottom; cluster-based permutation, 0.717–0.800 s, Σ*z* = 15.7, *p* = 0.029). This reduction was primarily driven by increases in the on-target fraction during wrong trials, rather than changes in correct trials, suggesting that after the switch, animals largely failed to show the increased exploration of alternative choices through off-target saccades in wrong trials that characterised pre-switch errors. Together with the post-switch lengthening of inter-saccadic intervals, this points to a scenario where after a rule switch, animals approach target choices that were formerly correct but are now wrong in a way that behaviourally resembles correct trials.

Saccadic entropy also shifted across the switch, but in an outcome-dependent manner (Fig. 5G; session-level Mann–Whitney): on correct trials, entropy decreased significantly (0.134 →0.121 bits, *p* = 0.022, *q* = 0.045, *r* = +0.36), whereas on wrong trials it increased but not significantly (0.290 →0.397 bits, *p* = 0.27, *r* = −0.18). The post-switch entropy change on wrong trials is therefore suggestive rather than established.

Whereas the lengthening of fixation durations was significant and uniform across outcomes (Fig. 5D), the entropy changes were smaller and outcome-dependent, with a significant decrease on correct trials and a non-significant increase on wrong trials. This suggests that the global slowing of saccadic timing and any outcome-specific change in gaze organisation are at least partly separable.

Together, these results dissociate three components of gaze across the switch. Target-directed orienting persisted, independent of stimulus identity. Inter-saccadic intervals lengthened uniformly, signalling the higher processing demand of the harder post-switch environment. Outcome-specific structure reorganised: the correct-versus-wrong divergence collapsed and entropy shifted, reflecting uncertainty about the new contingencies. Demand and uncertainty thus left separable signatures in gaze, while the underlying orienting strategy held constant.

## Discussion

In this study, we show that saccade dynamics in mice performing a visually guided foraging task are not simply stimulus-driven or reflexive, but reflect, among others, spatial predictions maintained in working memory, predicted trial outcomes, locomotor-oculomotor coupling, task difficulty and environmental variability, akin to the dynamics observed in more vision-centric mammalian species like primates and humans.

One should note that our findings are based on a relatively low acquisition frame rate of 60 Hz. Whilst most studies conducting analyses of oculomotor dynamics record saccadic eye movements at frame rates between 240 and 1000 Hz, our analyses focus on saccade occurrence, direction, and rate rather than on kinematic parameters such as peak velocity or precise duration, which require higher temporal resolution. Mouse saccades are large-amplitude, high-velocity movements (13), producing displacements that are robustly detectable as frame-to-frame position changes at 60 Hz even though their duration is short.

Furthermore, due to the limited availability of calibrated gaze-tracking systems for rodent research, as well as the added complication of using a spherical projection of the visual environment rather than a flat screen, we were not able to reliably extract exact visual gaze targets. As such, the current work focuses simply on the broad direction of saccades, but future work could further explore the target locations of saccades and eye fixations in mice.

Notably, although post-saccade running speed differed statistically from baseline, the difference was negligible in magnitude, and saccade deployment was not tightly coupled to locomotion. While locomotion is known to modulate activity throughout the mouse early visual system (34), this neural modulation was not strongly mirrored at the level of saccade deployment in our task. Instead, the task elicited distinct behavioural profiles across trial outcomes, including differences in path efficiency and reaction time as well as saccade patterns. In other words, animals adopted separable cognitive strategies that shaped their approach to each trial and ultimately determined its outcome, and these differential behavioural strategies included distinct saccading behaviour. Critically, such differential saccadic recruitment cannot be explained by stimulus-driven reflexive gaze shifts alone: animals presented with the same visual information deployed saccades differently depending on trial outcome, indicating that saccade generation was governed by internal decision state rather than by environmental statistics. This task-dependent modulation positions mouse saccades as a functional component of active visual sensing, consistent with the framework established in primates and contrary to prevailing assumptions about rodent saccades as reflexive oculomotor behaviour.

These findings stand in contrast with previous work showing saccade patterns in mice being largely shaped by low-level parameters like running speed, heading direction and gaze stabilisation (15, 17). This discrepancy may largely stem from a fundamental difference in experimental setup: in the absence of an immersive visual environment, coupled with naturalistic stimuli that the animals can explore and navigate around, locomotor activity might be the principal component driving saccades. In contrast, when the visual environment is dynamic, informative and task-relevant, saccades may be driven by broader cognitive and perceptual demands and strategies. Our study does not directly compare these contexts, and disentangling the contributions of visual engagement and locomotion would require within-subject manipulations of task demand. Nonetheless, these findings support the idea that mouse saccadic behaviour under visually demanding conditions is governed by task-relevant cognitive processes rather than low-level sensorimotor reflexes, and that the apparent simplicity of mouse saccades in prior studies may have been a product of the experimental context rather than a general species-level constraint.

Saccadic probability increased as a function of the animal’s lateral offset from its eventual decision target, indicating that saccades were preferentially recruited during moments of spatial discrepancy between the animal’s trajectory and its decision target. Saccade direction reliably predicted subsequent changes in running direction: a leftward saccade was typically followed by a leftward adjustment in trajectory, and vice versa. While this pattern is somewhat in line with the idea that saccades prepare the system for changes in heading direction (17), our results again do not point to a low-level sensorimotor reflex. Rather, saccade direction reliably predicted subsequent running direction changes across all trial outcomes, but most strongly on correct trials, where turns following leftward and rightward saccades were largest and most separable. This outcome-graded coupling between oculomotor and locomotor output points to a more flexible, higher-order role of saccades in navigational decision making rather than a fixed role in motor preparation.

In line with this notion, the temporal relationship between saccades and running direction was not fixed but varied systematically with the animal’s positional context. Larger lateral offsets were associated with pre-saccadic corrections, whereas smaller offsets were associated with post-saccadic corrections. This bidirectional, context-dependent timing extends previous observations that eye movements covary with turn direction during head-fixed plus-maze navigation (17): rather than a fixed temporal order between saccades and turns, our data point to a common underlying decision signal that coordinates oculomotor and locomotor outputs according to the urgency of the required correction. The fact that saccades and locomotor adjustments are temporally coupled, directionally congruent, and jointly sensitive to positional context establishes them as likely co-outputs of a shared decision process.

Saccades directed toward the eventual stimulus of choice (on-target saccades) emerged specifically after the onset of visual information that enabled discrimination, indicating that target-directed gaze shifts were contingent on the presence of actionable sensory evidence rather than being generated indiscriminately throughout the trial. Most strikingly, when animals retained prior knowledge of stimulus position, such as on post-error repeat trials where the stimulus location could be inferred from the previous trial’s outcome, they generated saccades toward their eventual choice before the stimulus was visually available. This anticipatory gaze direction dissociates saccade generation from immediate sensory input, indicating that mouse saccades can be driven by internally held spatial representations consistent with working memory. Critically, even these working-memory-driven saccades showed distinct coupling to trial outcome, demonstrating that state-dependent modulation persists independently of whether salient stimuli are present or not. Such non-stimulus driven saccades underline the importance of saccade strategy for visually guided success in our task structure. These results also point to a potential use of saccade patterns in future paradigms as a behavioural readout reflecting the uptake and maintenance of relevant information on a trial by trial basis even in the absence of external stimuli.

The temporal structure of saccadic behaviour, measured as saccadic entropy, was lower in correct than in wrong trials following stimulus onset, indicating a more ordered pattern of gaze deployment during successful performance. Rather than reflecting a fixed gaze sequence required to extract visual information, this structure points to differential saccadic recruitment governed by cognitive state. When spatial contingencies were already known, animals still exhibited low-entropy, on-target saccade recruitment, suggesting this structure reflects decision confidence rather than informational need. Further specificity emerged in the lateralisation effect described earlier: the preferential recruitment of saccades when the animal was spatially offset from its target was far stronger in correct trials and specific to saccades directed toward the eventual target. This dissociation argues against explanations based on general motor activation and instead implicates a targeted, cognitively governed sampling strategy. Taken together, these results indicate that the cognitive state of the animal is legible in the structure of its saccadic behaviour, and that this structure has functional consequences for performance. This further highlights the possibility of using oculomotor behaviour in mice as a non-invasive assay of internal representations, including highly accessible metrics like saccade entropy and spatial concentration of saccades.

A critical test of whether saccade patterns reflect a genuine cognitive process, rather than a learned sensorimotor habit tuned to a predictable environment, is how they are affected when environmental contingencies change. To characterise saccades in such variable contexts, we trained animals to switch between two stimulus sets with opposing behavioural contingencies.

That behaviour stabilises so rapidly across the switch is itself notable. Because the task is intuitive and naturalistic, animals adjusted to the reversed contingency within only one or two trials, allowing short contingency blocks and many switches within a single session, and pointing to a striking degree of behavioural flexibility. Overall performance remained comparable across the switch (Fig. 5B) even as the underlying cognitive demand increased, a dissociation we return to below, where the elevated demand becomes legible not in accuracy but in the timing of saccades and the directness of approach trajectories.

Following the introduction of rule switches, saccadic behaviour in wrong trials began to intermittently resemble that in correct trials, suggesting that saccade dynamics followed the task outcomes initially predicted by the animal. Saccadic entropy also shifted across the switch, but in an outcome-dependent manner: it decreased significantly on correct trials, while the increase on wrong trials did not reach significance (Fig. 5G). If saccadic entropy is taken to index the predictability of the animal’s gaze strategy, the significant reduction on correct trials is consistent with gaze becoming more stereotyped as the new contingency is consolidated. The tendency toward higher entropy on wrong trials, although not statistically reliable here, would be consistent with the possibility that post-switch errors reflect genuine uncertainty about which stimulus is currently rewarded rather than confident responding under a now-invalid rule (see (27, 35) for a related phenomenon); we note this interpretation as a hypothesis to be tested rather than an established result.

Two further signatures accompanied the rule switch and point in the same direction. First, inter-saccadic intervals lengthened significantly across correct and wrong trials (Fig. 5D), accompanied by a corresponding decline in saccade rate. Second, path efficiency decreased across the switch on both outcomes, with approaches becoming less direct (Fig. 5C). Together, a slowing of saccadic sampling and a reduction in the directness of approach trajectories are both consistent with the increased processing demand of the post-switch environment, even as overall accuracy was preserved. This global, outcome-independent lengthening of fixation durations is consistent with a demand-driven modulation of saccadic timing: a slowing of the sampling rhythm that tracks the overall processing load of the environment rather than the accuracy or strategy of any individual trial. A comparable demand-sensitivity of saccade rate has been described in human and macaque observers, where the rate of saccadic sampling is modulated by task and stimulus demands (31, 32).

Notably, in our task the visual scene was essentially unchanged across the switch: the same stimuli were presented, and only their behavioural significance became uncertain. The lengthening of inter-saccadic intervals is therefore unlikely to reflect increased visual processing load, and instead points to a cognitive contribution to saccadic timing. From this perspective, mice occupy the slow end of a sampling continuum that, while differing several-fold in absolute timescale from primates, is similarly demand-sensitive, rather than reflecting a categorically distinct mechanism. This suggests that the additive component pacing each fixation is not specifically visual, but reflects total processing demand, to which bottom-up visual complexity and top-down cognitive difficulty contribute through a common channel.

These data thus point to two partly separable scales of saccadic modulation: a global timing scale, on which fixation duration lengthens significantly with processing demand in a manner qualitatively consistent with the demand-sensitive modulation of saccade rate described in primates (32), and a cognitive scale, on which orienting strategy and saccadic entropy shift after the switch in an outcome-dependent way. That both are legible in the same trials, with the cognitive modulation superimposed over rather than replacing the demand-driven baseline, is consistent with a control architecture in which sensory and cognitive demands jointly shape oculomotor output.

Our findings demonstrate that mice, when performing a task that demands active visual engagement, deploy saccades as part of a cognitively modulated, decision-relevant active sampling strategy. The classical view that mice lack the retinal and behavioural infrastructure for active visual sensing may have been too quick to equate the absence of a classical retinal fovea with the absence of visual sampling strategies, and the reliance on non-visual senses with the redundancy of vision. In contrast to this notion, our data show that the relevant question was never whether mice possess the capacity for cognitively driven saccadic behaviour, but whether they had been given a reason to use it. Mice may saccade for different functional reasons than primates, perhaps to modulate visual input gain rather than to fixate a foveal target; consistent with such a role, mouse V1 distinguishes the visual motion induced by the animal’s own saccades from externally generated motion, indicating that saccades are treated as functionally significant events by the early visual system (36). Yet the state-dependent structure of their saccadic behaviour nonetheless echoes what is observed in primates. That this capacity parallels primate saccadic dynamics in its temporal structure, state-dependence, and sensitivity to cognitive load suggests that active visual sensing may be a more broadly conserved strategy across species than current models assume. These findings open the door to using mouse oculo-motor behaviour as a non-invasive, high-temporal-resolution readout of cognitive state, decision confidence, and working memory, capitalising on the same genetic, imaging, and electrophysiological tools that make the mouse an indispensable model system, now applied to a behavioural domain previously thought to be beyond its reach.

## Acknowledgments

We thank Olga Arne for assistance with the experiments, and Dr Katharine Shapcott and Dr Marvin Weigand for their help in setting up DomeVR. We thank Dr Abdel Nemri and Dr Tim Schröder for help with hardware development and setup. We thank the Ernst Strüngmann Institute Animal Facility, including the animal caretakers and veterinarians, for supporting our work in animal husbandry and care. We also thank the Ernst Strüngmann Institute core facility and IT teams for supporting experimentation and computational infrastructure. Finally, we acknowledge the animals who made this work possible. This work was supported by the Max Planck Society.

## Author Contributions

Conceptualization, R.T., M.S. and M.H.; Methodology, R.T.; Software, R.T., C.W., M.Y.A.E.H., and M.G.; Validation, R.T. and M.Y.A.E.H.; Formal Analysis, R.T.; Investigation, R.T.; Resources, R.T., M.S. and M.H.; Data Curation, R.T. and M.Y.A.E.H.; Writing – Original Draft, R.T.; Writing – Review & Editing, R.T., M.H., and M.S.; Visualization, R.T.; Supervision, M.H. and M.S.; Project Administration, M.H. and M.S.; Funding Acquisition, M.H. and M.S.

## Declaration of Interests

The authors declare no competing interests.

**ACKNOWLEDGEMENTS**

## STAR Methods

### Key Resources Table. Resource Availability

#### Lead contact

Further information and requests for resources should be directed to and will be fulfilled by the lead contact, Robert Taylor.

#### Materials availability

This study did not generate new unique reagents.

#### Data and code availability

- All behavioural and eye-tracking data reported in this paper have been deposited at Zenodo and are publicly available as of the date of publication. The DOI is listed in the key resources table.
- All original code has been deposited at Zenodo and is publicly available as of the date of publication. The DOI is listed in the key resources table.
- Any additional information required to reanalyse the data reported in this paper is available from the lead contact upon request.

### Subjects and surgical procedures

#### Animals

All procedures were performed in accordance with the German law for the protection of animals and the European Union’s Directive 2010/63/EU (authorisation number F149/2000, *Regierungspräsidium Darmstadt*). Eleven male FVB/NRj × C57BL/6NRj F1 mice (bred in-house from Janvier Labs parental strains) were used, with ages at the start of experimentation ranging between 8 and 25 weeks. Mice were kept on a reversed day–night cycle and were initially group-housed, but housed individually once behavioural training commenced to ensure reproducible and safe food restriction, as well as to prevent damage to neural implants and head-plates. Single-housed animals were given regular ‘playtime’ in a large play cage shared with litter mates after task training sessions in order to counteract stressful effects of single housing. Mouse cages were placed in a room with a temperature of 21.5 °C and 55% humidity, and contained environmental enrichment including running wheels, wooden toys, and shelters. Mice were handled extensively for 5 days prior to the initiation of behavioural training. Data collection, handling, and husbandry were conducted exclusively by the first author to maintain familiarity.

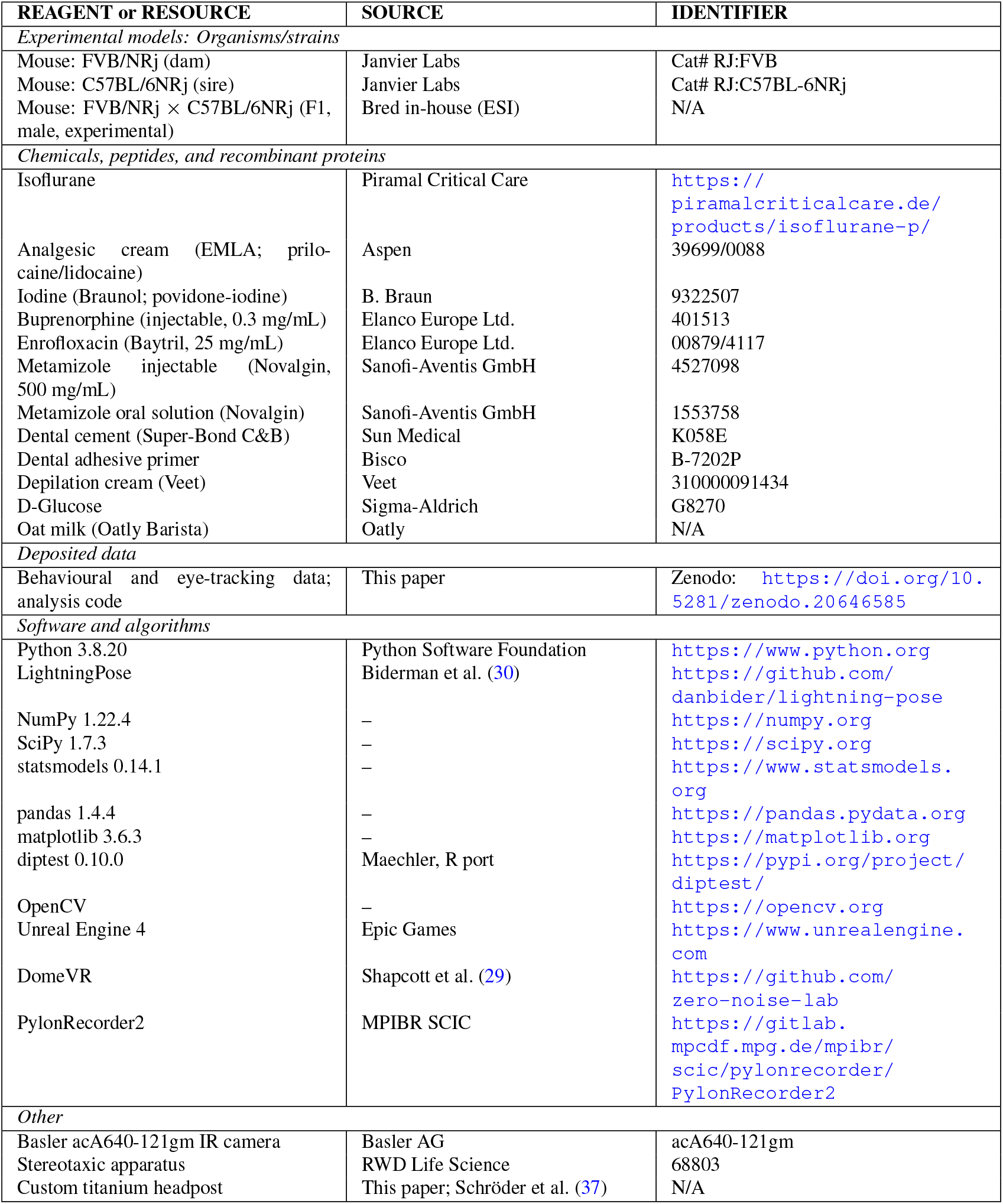

#### Surgical procedures

Mice were fitted with custom-milled titanium headposts for the duration of the experiment (37). Animals were placed on a stereotaxic apparatus (RWD 68803) under isoflurane anaesthesia (induction 3–4%, maintenance 1–2% in O_2_). The surgical site was shaved and depilated with depilation cream (Veet), and the skin prepped with povidone-iodine (Braunol). Lidocaine was injected subcutaneously at the incision site for local analgesia, and EMLA cream (prilocaine/lidocaine) was applied topically. An incision was made and the skin on top of the cranium was removed; the cranium was then cleaned, treated with a dental adhesive primer (Bisco B-7202P), and the custom-milled titanium head plate attached using dental cement (Super-Bond C&B, Sun Medical). For post-operative analgesia and antibiotic prophylaxis, mice received subcutaneous buprenorphine (0.1 mg/kg) and injectable enrofloxacin (20 mg/kg), followed by enrofloxacin (0.17 mg/mL) and metamizole (1.25 mg/mL) in the drinking water for 3 and 5 days post-surgery, respectively; depending on water intake, mice received approximately 1.02 mg/day of enrofloxacin and 7.5 mg/day of metamizole. Mice were given a minimum of 5 days to recover before the initiation of handling and behavioural training. For full details on surgical procedures, see Schröder, Taylor et al. (37).

#### Food restriction

Animals were kept lightly food-restricted for the duration of data collection. At the start of handling, food rations were gradually reduced from 4–5 g of dry food (typical daily intake for mice of this sex and age) to 2.5–4 g, allowing for the stable maintenance of a body weight between 90–95% of expected body weight from an individually fitted growth curve adjusted for age. Food was administered 2–4 h after the completion of the task session, which typically fell 2–4 h before the end of the animal’s dark cycle.

### Experimental Setup

#### Virtual reality apparatus

Mice were head-fixed using a custom-built head-fixation mechanism (37) and placed on a styrofoam trackball allowing unconstrained 2D locomotion within a virtual environment. The virtual environment was designed in Unreal Engine 4, allowing for the precise timing and control necessary for neuroscience experiments, and projected onto a 250° dome-shaped enclosure via a curved mirror at 60 Hz (see Shapcott et al. (29) for details). Mice were placed at the centre of the 120 cm diameter dome onto which the virtual environment was projected, covering both the central and peripheral visual field within a dynamic naturalistic environment the animal could actively traverse.

#### General task design

Our task was based on a virtual reality foraging paradigm in which mice navigated freely within a naturalistic virtual environment, with unrestricted movement along all rotational axes and translational movement in two dimensions. Animals were required to navigate towards salient rewarded stimuli whilst ignoring unrewarded distractors. Upon contact with a rewarded virtual stimulus, mice automatically received a drop of oat milk with 5% glucose dissolved, delivered via a lick spout. The virtual arena consisted of dynamically responsive grass in the fore-ground, with mountains in the background and a blue sky with dynamic clouds overhead. Stimuli were modelled on leaf shapes with naturalistic leaf textures, providing randomness and ecologically relevant visual properties.

##### Post-error trials

Following an incorrect or missed response, the same stimulus configuration was re-presented so that animals did not have the option to avoid visually difficult trials by completing them inaccurately. This also meant that in repeat trials, the location of the rewarded stimulus was in principle predictable.

#### Task training

Training proceeded through a series of stages designed to incrementally shape the animals’ behaviour. In Stage 1, the animal’s lateral movement was restricted by hedge corridors flanking the path, constraining the mouse to run straight towards the rewarded stimulus and thereby establishing the association between the stimulus and the reward. This stage lasted between 1–3 training sessions. In Stage 2, the hedges were removed, requiring the animals to learn to execute lateral turns whilst maintaining stable, straight running trajectories. This stage lasted 1–5 sessions. Once animals demonstrated proficient straight running, lateral offsets of the stimulus from the centre of the arena were gradually introduced, training the animals to deviate from straight paths in order to obtain the reward.

Stage 3 introduced an ambiguous placeholder stimulus that displayed the natural statistics of both the rewarded stimulus and the later-introduced distractor. Animals were trained to run towards this placeholder, which, on their approach, moved laterally (left or right) and away from the animal, matching the structure of the final task. The distractor was then gradually introduced, diverging from the same ambiguous placeholder. Initially, the distractor appeared at very high transparency to minimise novelty bias, and its opacity was gradually decreased over subsequent sessions until it matched that of the rewarded stimulus. At this point, animals had reached the final discrimination stage, which is the stage included in the present analysis. The duration of this final training stage varied between 3–10 sessions.

##### Basic discrimination training

Each trial began with an ambiguous placeholder stimulus presented directly ahead of the animal. The animal was required to navigate toward the ambiguous placeholder to initiate the trial, at which point the placeholder divided into a rewarded stimulus (S+) and an un-rewarded distractor (S −) placed at a lateral offset and spawning randomly on either the left or right side of the arena. The stimuli were objects with leaf-like texture. The S+ outline was of a compound palmate leaf with 7 broad lanceolate leaflets, culminating in a point. The S− outline consisted of two rounded lobules, one placed over the other (Fig. 1B). The animal navigated toward one of the two stimuli within arena space to register a choice. Correct responses (contact with S+) were rewarded. Incorrect responses (contact with S−) triggered a timeout period during which the animal was teleported into a penalty corridor, and had to navigate out. Missed trials (failure to hit either stimulus) caused the trial to restart.

##### Generalisation training

To promote stimulus generalisation, leaf shapes created by morphing the S+ and S− in different proportions were introduced in 12 successive stages. First the S+ and then the S− were each gradually replaced by stimuli geometrically closer to a 50% morph (the geometric midpoint between the original S+ and S−), such that the animal learned to treat the same 50% morph as either S+ or S− depending on the identity of the paired stimulus.

##### Rule context switch

After generalisation training, the role of the 50% morph alternated between S+ and S− in uncued blocks of 10 correctly responded trials (5–20 trials in total, median 12). Mean number of switches in a single session was 27.0 ± 10.3. Only animals that had reached a stable, high level of behavioural performance during generalisation training progressed to the switch condition. This was assessed qualitatively by the experimenter rather than against a fixed numerical criterion, on the basis of two indicators: consistent, high accuracy across sessions, and stable path efficiency (reflecting direct, non-exploratory approaches to the chosen stimulus).

#### Randomisation and blinding

Stimulus side (left vs. right of the arena midline) was randomised on each trial by the virtual reality software. No experimenter blinding was applied, as all task contingency manipulations were automated by the VR software and were within-subject, driven by the animal’s own behaviour rather than by experimenter assignment.

#### Eye tracking

Bilateral pupil position was recorded at 60 Hz using two Basler acA640-121gm infra-red cameras with a modified version of PylonRecorder2 software (https://gitlab.mpcdf.mpg.de/mpibr/scic/pylonrecorder/PylonRecorder2). To synchronise the video timing with events in the virtual reality environment, we used 32 ms long infra-red flashes emitted from an LED mounted near the camera lens at the onset of each trial. These flashes were then extracted from the videos to be used as timestamps for synchronisation with DomeVR. Five consecutive flashes indicated the start of a behavioural session; a single flash indicated the start of a trial. To select sessions with reliable pupil estimates, the per-frame, per-keypoint LightningPose confidence scores (range 0–1) were averaged across all frames and keypoints within each session, and sessions with a mean confidence above 0.97 were retained. This criterion yielded 140 of 205 sessions across all 11 animals.

Pupil position was extracted offline using LightningPose via markerless pose tracking. A single model was trained across animals and sessions: frames were sampled from a wide array of animals and sessions, and labelled with 8 keypoints on the circumference of the pupil. For each frame, the (*x, y*) coordinates of all LightningPose-tracked keypoints surrounding the pupil were extracted. Keypoints were excluded on a per-frame basis if their likelihood fell below 0.99 or if their summed coordinates fell outside the 0.05th–99.95th percentile range of the session distribution, to remove low-confidence and outlier detections. An ellipse was then fitted to the remaining valid keypoints (minimum 5 points required) using OpenCV’s direct least-squares ellipse fitting algorithm, and the centre of the fitted ellipse was taken as the pupil position for that frame. Frames with fewer than 5 valid keypoints were assigned missing values and excluded from subsequent analysis. Ellipse centres were z-scored to the session mean and standard deviation, removing between-session and between-eye differences in absolute pixel position and scale, derived from changing crop sizes, camera distances, and between-animal variability. We restricted analyses to a single eye after confirming highly symmetric conjugate eye movements across both eyes (Fig. S2).

#### Saccade detection

Frames coinciding with camera synchronisation flashes were blanked (NaN for 3 frames post-flash). The z-scored horizontal trace was smoothed with a Savitzky– Golay filter (window = 15 frames, polynomial order = 1), and frame-to-frame velocities were computed as first differences. Velocity samples following *≥* 4 consecutive missing frames were suppressed for 5 frames to prevent interpolation artefacts. Saccades were detected using an adaptive median absolute deviation (MAD) based threshold: samples with velocities exceeding ± 5 MAD from the session median were identified, grouped into contiguous same-sign runs, and retained if spanning *≥* 5 frames of the smoothed z-scored horizontal ellipse centre trace. Runs of the same direction within a *≤* 10 frame window in the smoothed trace were merged. Saccade onset was defined as the first frame of each detected run. Per-saccade amplitude was computed from the raw z-scored trace over the detected saccade window, normalised by frame dimensions (i.e., expressed in units of fractional frame width rather than degrees of visual angle). All traces were segmented into per-trial epochs aligned to flash-derived trial boundaries.

#### Behavioural metrics

*Correct, wrong, and miss outcomes* were defined as hitting the correct stimulus, hitting the wrong distractor, and missing both targets entirely, respectively. Cohort-level outcome proportions (Fig. 1C) were computed across all task trials, including post-error repeat trials, with probe trials excluded.

*Hit rate* was defined as the percentage of correct trial outcomes within a given session. Because the open-arena task does not force a binary choice and requires animals to actively navigate toward a stimulus, the uninformed success rate is very low: in a closely related virtual-environment task, removing visual input abolished directed responses entirely (28).

*Path readout* was collected as a list of *x* and *y* coordinates within arena space per refresh of the virtual environment (60 Hz); see Shapcott et al. (29). Arena distances at a given translation ratio were then converted to cm by measuring the distance the trackball travelled per unit within the arena.

*Path efficiency* was defined as the ratio of the optimal to the actual path length:

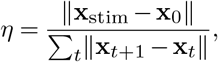

where **x**_0_ is the animal’s position at stimulus onset, **x**_stim_ the centre of the chosen stimulus, and the denominator is the cumulative path length of the actual trajectory. Values approach 1 for direct approaches and decrease with detours.

*Reaction time* was computed from the *y*-axis displacement of each trial’s trajectory within arena space. Raw position data were first baseline-corrected by subtracting the initial coordinates, then low-pass filtered using a first-order Butterworth filter (cutoff = 3 Hz, sampling rate = 60 Hz). A sliding-window *R*^2^ approach was used to detect movement onset: linear regression was fit across rolling windows of multiple sizes, and *R*^2^ scores were computed for each window position. Drops in *R*^2^ (indicating deviation from linearity, i.e., movement initiation) were identified as peaks in the inverted *R*^2^ signal. A normalised Gaussian kernel was applied at each detected peak to produce a continuous weight array. Weight arrays from all window sizes were summed and multiplied by a linear time-decay function (1 to − 0.1) to bias detection toward earlier timepoints. Reaction time was defined as the sample index corresponding to the maximum of this combined weight signal. Trials with insufficient data points or anomalously low peak weights (below the 5th percentile of the per-session distribution) were excluded.

*Running speed* was computed from the trackball readout as Euclidean displacement per frame in arena space. This was then converted into cm s^−1^ by translating arena distance units into rotational distance of the styrofoam trackball at the given translation factor.

#### Saccade analysis

##### On-target classification

Saccades were classified as on-target or off-target relative to the chosen stimulus, defined as the stimulus the animal ended up hitting at the conclusion of a given trial, regardless of stimulus identity. Saccades were classified by comparing the sign of the horizontal saccade direction to the sign of the animal’s lateral offset from the chosen stimulus at saccade onset; a saccade was scored as toward when these matched and away otherwise. Because the eye-camera and arena reference frames are not jointly calibrated, this convention was validated empirically: on trials with a strongly lateralised target that the animal demonstrably approached, the fraction of saccades classified as toward the chosen stimulus exceeded chance both across animals and across pooled saccades (Fig. 3E; see Results), confirming that the classification reflects gaze directed toward, rather than away from, the chosen stimulus.

##### Fraction on-target over time

The fraction of on-target saccades was computed in time bins of 16.7 ms aligned to stimulus onset, separately for correct and wrong trials. Shaded regions are descriptive, with statistical inference provided by the cluster-based permutation test described below.

##### Saccade rate

Saccade rate (Hz) was computed as *N*_saccades_*/T*_post_, the number of post-stimulus saccade onsets divided by the post-stimulus trial duration (s).

##### Inter-saccadic intervals

Inter-saccadic intervals (ISIs) were computed as the time between successive saccade onsets. For the rule-switch analysis (Fig. 5D), full-trial ISIs on stimulus-present correct and wrong trials were compared between pre- and post-switch conditions. Pre-switch data comprised the non-switch sessions immediately preceding each animal’s first switch session; post-switch data comprised the switch trials together with their subsequent trials within each switch session. ISIs were pooled across trials within each condition (*n* = 1,770 pre-switch / 961 post-switch on correct trials; 1,330 pre-switch / 614 post-switch on wrong trials) and compared using Mann–Whitney *U* tests, separately for correct and wrong trials (see Quantification and Statistical Analysis).

##### Saccade amplitude tertiles

For analyses requiring amplitude-binned saccades (Fig. S3), saccades were split into tertiles based on the within-session amplitude distribution. Tertile cutoffs were therefore computed dynamically per session rather than using a fixed threshold across the dataset.

##### Turn-change angle

The angular change in heading direction was computed as the difference between the mean heading over the ± 30 frames (0.5 s) preceding saccade onset and the mean heading over the 60 frames (1.0 s) following it.

##### Turn latency

The latency of maximal turn amplitude was defined as the time of peak absolute heading change velocity relative to saccade onset. This was computed at each candidate timepoint *t* in the search window of ±30 frames around a saccade onset. Mean displacement was calculated in the pre-segment of 5 frames before and post-segment of 5 frames after *t*. Angular difference was then computed between each pre- and post-segment at *t*, and the *t* with the largest angular difference was taken as the latency relative to saccade onset. *Saccade entropy*. Per-trial entropy was computed as the binary Shannon entropy of the on-target saccade proportion:

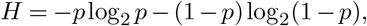

where *p* is the fraction of saccades directed toward the chosen stimulus, counted over the post-stimulus window (from stimulus onset to the end of the trial epoch). To avoid singularities, *p* was clamped to [10^−6^, 1 − 10^−6^]. Trials with no post-stimulus saccades were excluded; trials with at least one saccade were retained (single-saccade trials therefore take *H ≈* 0). This measure captures the predictability of saccadic allocation on each trial, yielding maximum entropy when gaze is divided equally between stimuli and minimum entropy when directed exclusively toward one option. Because per-trial *H* was averaged across trials before comparison, the resulting measure is sensitive to trial-to-trial consistency rather than to the pooled proportion alone: a condition in which gaze is decisively directed on every trial yields low mean entropy, whereas the same overall on-target proportion achieved through a mixture of inconsistent trials yields higher mean entropy. Mean entropy therefore indexes the trial-to-trial reliability of oculomotor decision-making, complementing the pooled on-target fraction (Fig. 3C, E). For all entropy comparisons (Fig. 3D, Fig. 4D, Fig. 5G), per-trial *H* was averaged per session within each condition, and these per-session values were compared using Wilcoxon signed-rank tests (correct vs. wrong, Fig. 3D, Fig. 4D) or Mann–Whitney *U* tests (pre-vs. post-switch, Fig. 5G), with *n* denoting the number of sessions. The session-level summary was used because per-trial *H* is strongly zero-inflated (most trials contain a single in-window saccade, giving *H ≈* 0), which makes a per-trial test sensitive to the heavy upper tail rather than to the central tendency reflected in the figure.

##### Saccade onset density

Two-dimensional histograms of saccade onset position (*dx × dy*) were computed relative to the chosen stimulus, using a bin size of 6.67 arena units per bin (forward) and 10 arena units per bin (lateral). Colour scale shows log(1 +count).

##### Lateral density distributions

Density curves for the lateral component (*dy*) of saccade direction were estimated using histogram binning (120 bins) smoothed with a Gaussian kernel (*σ* = 1.4 bins).

#### Quantification and Statistical Analysis

All statistical tests were two-sided unless otherwise stated, and evaluated at *α* = 0.05. Data were collected from 11 animals across 140 of 205 sessions retained by quality control (mean left-eye LightningPose confidence > 0.97), with no animal excluded; nine animals reached the uncued rule-switch condition. Unless otherwise stated, comparisons of saccade- and trial-level measures (on-target proportion, inter-saccadic interval, path efficiency, turn angle, post-saccade running speed) were performed on observations pooled across animals, with *n* denoting the number of saccades or trials as specified for each test in the Results. Saccade entropy (Fig. 3D, Fig. 4D, Fig. 5G), and the saccade-rate comparison between outcomes (Fig. 3B), were instead compared at the session level, with each session contributing a single value and *n* denoting the number of sessions; the across-switch saccade-rate comparison (Fig. 5E) was performed on pooled trials. The comparison of overall task performance across the rule switch (Fig. 5B) was performed at the animal level (*n* = 9 animals), as it concerns a per-animal behavioural summary. Because pooled saccade- and trial-level tests treat correlated observations within animals as independent, the resulting *p*-values index the reliability of a difference in the pooled distribution rather than its magnitude; we therefore report an effect size alongside every test (see below) and base interpretation on effect size as well as significance. All statistical details (test, *n*, effect size, *p*, and where applicable *q*) are reported in the Results.

##### Multiple-comparison correction

To control the false discovery rate across related comparisons, Benjamini–Hochberg correction was applied within each family of related tests (for example, the set of pairwise outcome contrasts for a given metric, or the set of chance-level comparisons within a figure panel). Corrected values are reported as *q* alongside the uncorrected *p* in the Results; the family to which each correction applies is indicated where relevant. Single, standalone comparisons were not corrected. For the cluster-based permutation tests, multiplicity across the time axis is controlled within the test itself by the maximum-statistic null distribution (see below), and cluster *p*-values are therefore reported without further correction.

##### Cluster-based permutation tests

were used to identify sustained temporal differences in the on-target saccade fraction between correct and wrong trials (Fig. 3C, Fig. 4C, Fig. 5F). The on-target fraction was computed in 16.7 ms bins (60 Hz) aligned to stimulus onset. At each time point, on-target and off-target saccade counts were pooled across a centred 5-bin window (*∼*83 ms), and a two-sample *z*-test for equality of proportions between correct and wrong trials was computed on the windowed counts; the two bins at each edge of the time axis, where a full window could not be formed, were excluded, giving an effective range of ± 0.967 s. This moving-window pooling increases the per-time-point sample size relative to a single-bin test and was used in place of post-hoc smoothing of the *z*-trace. Clusters were defined as contiguous time points at which the absolute windowed *z* exceeded 1.96 (*α* = 0.05, two-tailed), with the cluster-level statistic the summed *z* within each cluster (Σ*z*; positive values indicate a higher on-target fraction on correct than wrong trials, negative the reverse). A null distribution was constructed by randomly shuffling trial outcome labels (correct versus wrong) 5,000 times and recomputing the entire windowed-test and clustering procedure on each permutation, retaining the maximum absolute cluster mass. Observed clusters were assigned *p*-values as the fraction of permutations with a larger maximum cluster mass; this maximum-statistic null controls family-wise error across the time axis, and cluster *p*-values are therefore reported without additional correction. The permutation used a fixed random seed for reproducibility.

##### Occupancy-normalised saccade location

For analyses of where saccades occurred as a function of lateral offset from the chosen stimulus (Fig. 2E, Fig. 3F), raw saccade-onset counts were divided by the dwell time (number of frames) the animal spent at each offset bin, yielding an onset rate (Hz) that controls for uneven occupancy across the offset axis. Bins with fewer than 50 dwell frames were masked. Bimodality of the occupancy-normalised offset distribution (Fig. 3F) was assessed with Hartigan’s dip test; results were stable across bin counts (40–80) and dwell thresholds (50– 200 frames). For Fig. 3F this normalisation was applied separately to on-target and off-target saccades, using a common per-bin dwell denominator so that the relative difference between classes reflects gaze allocation rather than occupancy. *Group comparisons*. Two-group comparisons of saccade- and trial-level measures (inter-saccadic interval, path efficiency, post-saccade running speed, turn angle, and the session-level saccade-rate and entropy comparisons) were performed using the Mann–Whitney *U* test for independent samples and the Wilcoxon signed-rank test for paired samples, as specified for each test in the Results. For these tests effect size is reported as the common-language effect size (CLES) and the rank-biserial correlation *r*. Proportions of on-target saccades were compared against chance (*p* = 0.5) using two-sided binomial tests, and between outcomes using *χ*^2^ tests of independence; effect sizes are reported as Cohen’s *g* for one-sample proportions and, for *χ*^2^ contingency tests, Cramér’s *V* together with the odds ratio for 2 × 2 tables. The per-animal comparison of task performance across the switch (Fig. 5B) used the Wilcoxon signed-rank test across animals, with effect size as the matched-pairs rank-biserial correlation *r*.

##### Circular statistics

Circular density plots (Figs. 2–5) were smoothed with a wrapped Gaussian kernel for visualisation but were not subjected to circular hypothesis tests.

##### Bimodality

Hartigan’s dip test was used to assess bimodality of lateral offset (*dy*) distributions (Figs. 3, 5), implemented via the Python diptest package v0.10.0 (a port of the R diptest package by Hartigan & Hartigan, 1985).

All analyses were performed in Python 3.8.20 using NumPy 1.22.4, SciPy 1.7.3, statsmodels 0.14.1, pandas 1.4.4, mat-plotlib 3.6.3, and diptest 0.10.0. No analyses were pre-registered.

## Supplementary figures

**Fig. S1.**
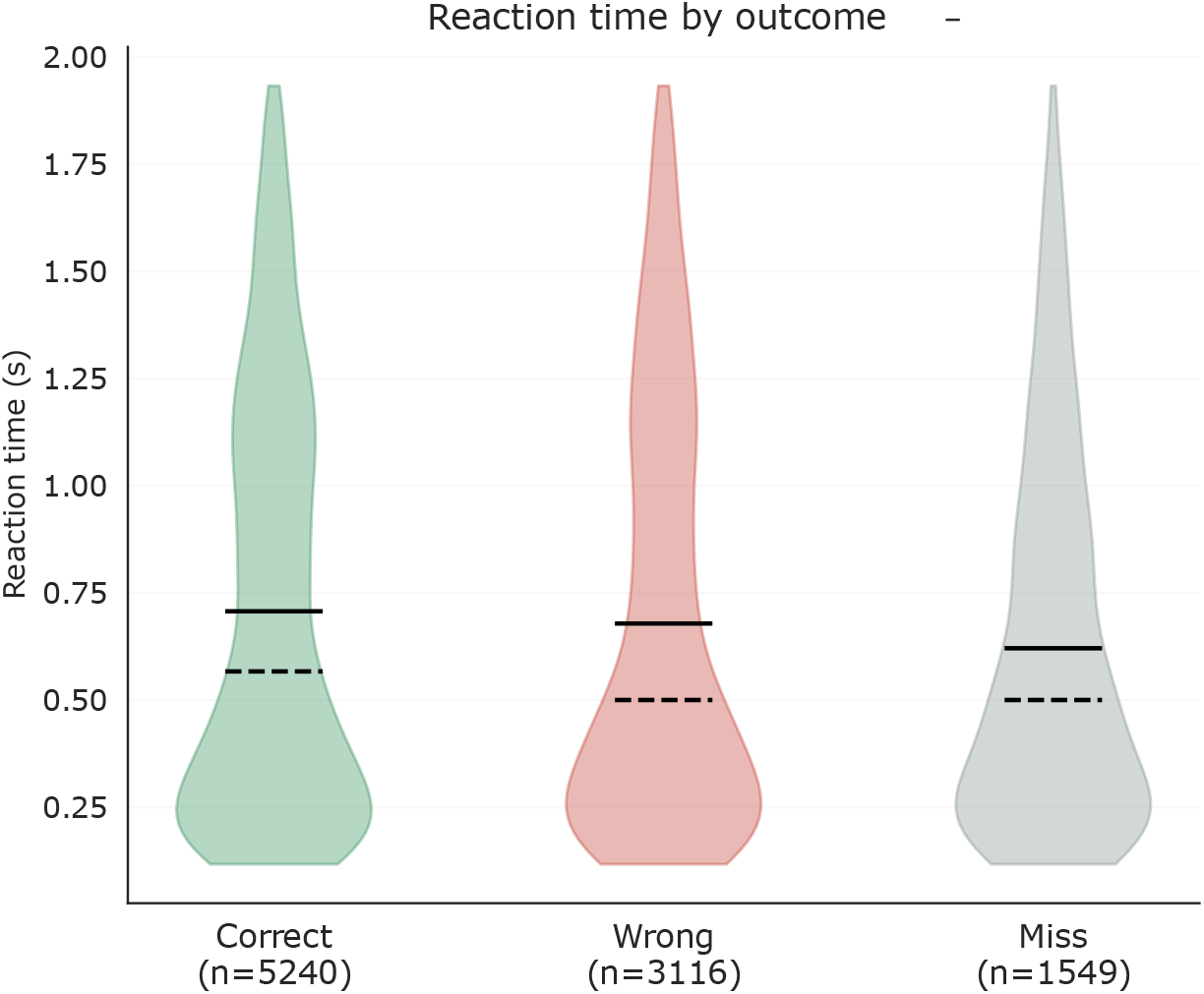
Reaction time distributions by trial outcome. Violin plots of reaction times (see Methods) for correct (green), wrong (red), and miss (grey) trials, pooled across all sessions and animals (n values indicate trial counts). Solid lines indicate the mean; dashed lines indicate the median. Reaction times were modestly longer on correct trials than on wrong or miss trials. Related to Fig. 1.

**Fig. S2.**
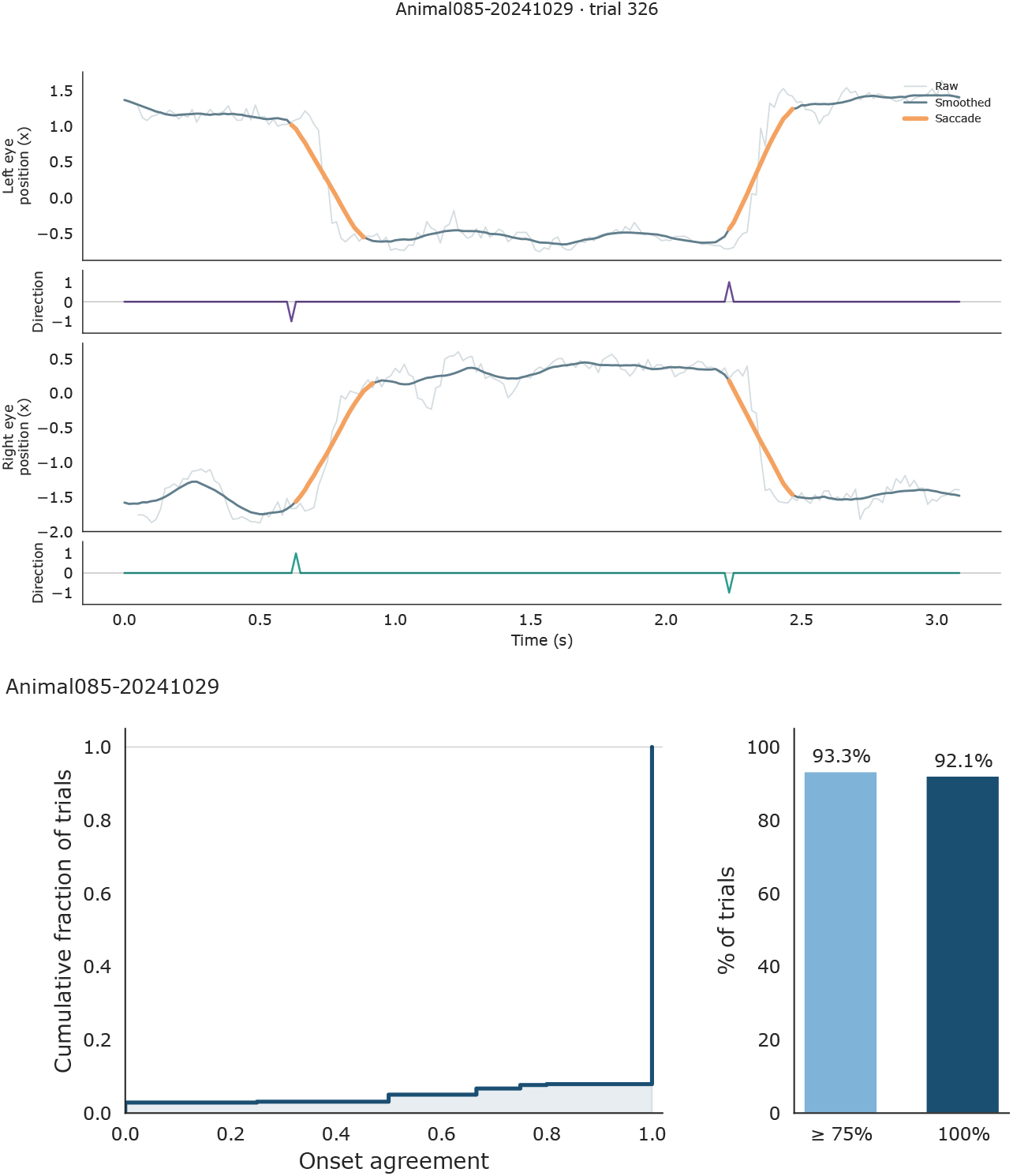
Horizontal eye movements are conjugate between the two eyes. To establish whether the two eyes move together during task performance, we analysed a single representative session with high-confidence bilateral pupil tracking (Animal085, session 2024-10-29), isolating the question of eye-movement conjugacy from variability in tracking quality. **(A)** Example trial (trial 326) showing horizontal pupil position for the left (top) and right (bottom) eye. Light grey: raw trace; dark teal: smoothed trace; orange: detected saccade epochs. Subpanels beneath each trace indicate detected saccade direction (±1). Saccades were detected independently in each eye and showed coincident events of opposing sign, consistent with conjugate horizontal movements. **(B)** Cumulative distribution of onset agreement across trials within the session, where onset agreement is the fraction of saccades per trial with a matched saccade in the contralateral eye within a 50 ms window (see Methods). **(C)** 93.3% of trials showed *≥* 75% onset agreement and 92.1% showed perfect (100%) agreement between the two eyes, indicating that tracking one eye is sufficient to capture binocular saccadic dynamics. Related to Fig. 1.

**Fig. S3.**
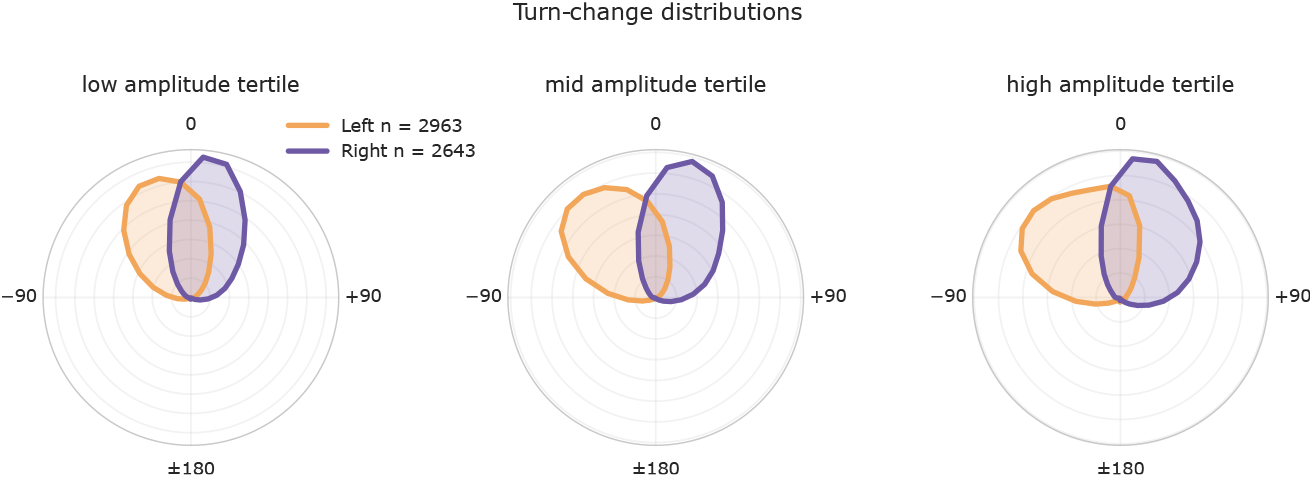
Saccade–turn coupling is preserved across saccade amplitudes. Polar distributions of the maximum running-direction change associated with each saccade (see Methods), split into tertiles of saccade amplitude (low, mid, high) and shown separately for leftward (orange, *n* = 2963) and rightward (purple, *n* = 2,643) saccades. 0° = no change, ±90° = left/right turn. Radial axis: probability density. Saccade–turn coupling is preserved across amplitudes. Related to Fig. 2.

**Fig. S4.**
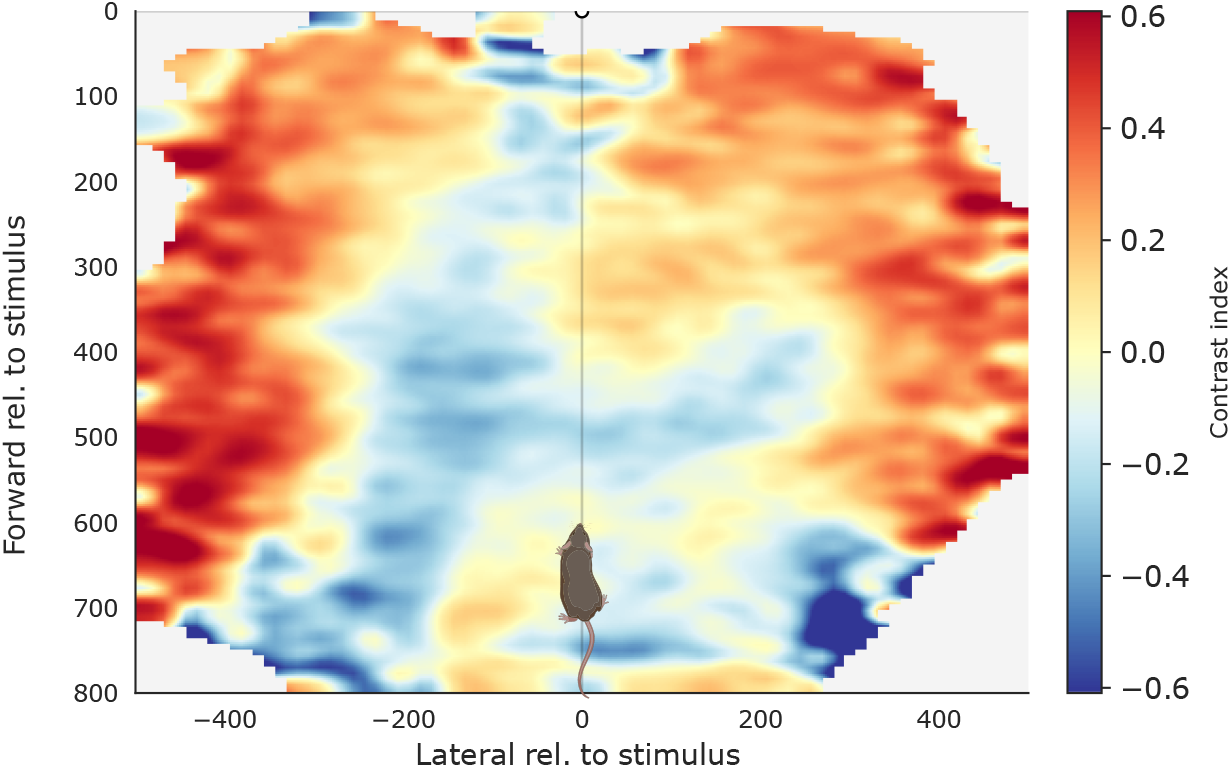
Saccades are spatially structured within the VR arena. Spatial distribution of saccade occurrence normalised by occupancy, expressed as a contrast index CI = (*P*_event_ − *P*_occupancy_)/(*P*_event_ + *P*_occupancy_), where *P*_event_ is the spatial probability of saccade onset and *P*_occupancy_ is the spatial probability of the animal’s position. Positive values (red) indicate regions where saccades occur more frequently than expected from occupancy alone; negative values (blue) indicate relative suppression. Coordinates are expressed in VR units relative to the stimulus location at (0, 0). The mouse silhouette marks the trial start location; the cannabis-leaf icons indicate the two possible stimulus locations (only one stimulus appeared per trial; see Methods). Data pooled across all animals, sessions, and trial outcomes. Related to Fig. 3.

**Fig. S5.**
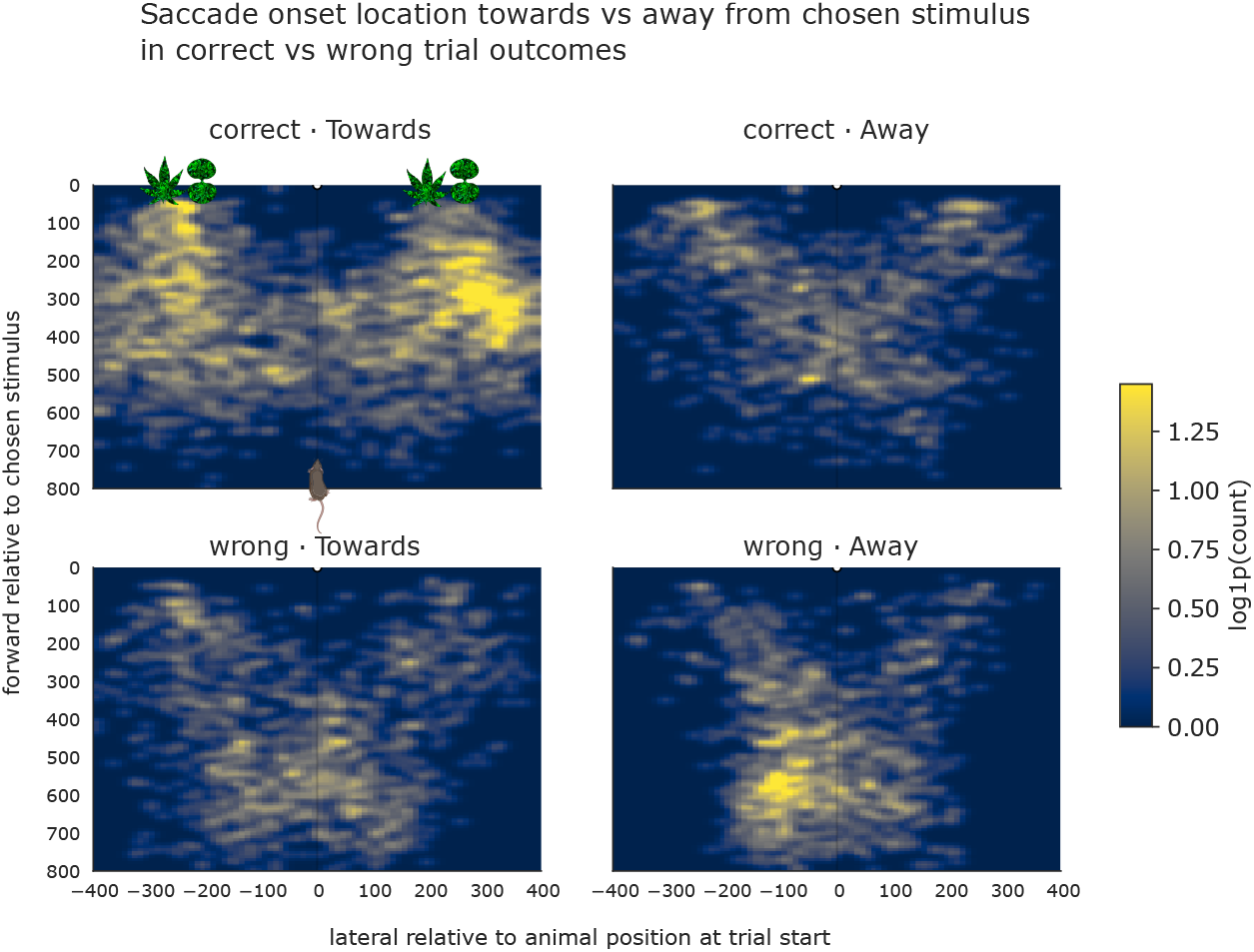
Spatial distribution of saccade onsets depends on saccade direction and trial outcome. 2D histograms of saccade onset locations in VR arena coordinates, split by trial outcome (rows: correct, wrong) and saccade direction relative to the chosen stimulus (columns: towards, away; see Section E). *x*-axis: lateral position relative to the animal’s trial-start position (VR units). *y*-axis: forward position relative to the chosen stimulus, with 0 at the stimulus (top) and higher values closer to the trial-start position (bottom). Stimulus icons (top-left panel) mark the two possible stimulus locations; mouse silhouette marks the trial-start position. Colour indicates log(1 + count) of saccades per spatial bin. Only post-stimulus saccades are included. Miss trials are excluded as no stimulus is chosen on miss trials and towards/away cannot be defined. Related to Fig. 3.

**Fig. S6.**
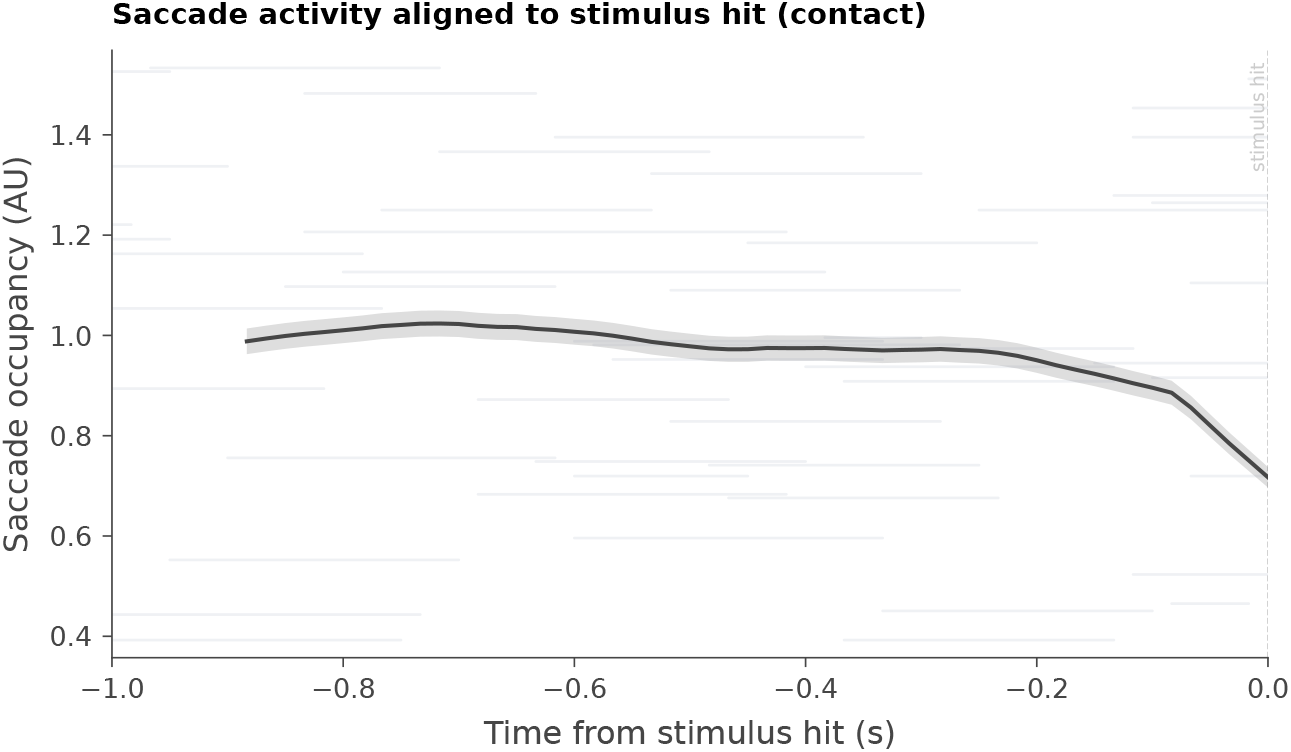
Peri-stimulus-hit saccade raster and PSTH. As in Fig. 1I, but with saccade events aligned to stimulus contact (the moment the animal reaches its chosen stimulus) rather than to stimulus onset. Grey: individual saccade events, each row depicting one trial; black: population moving sum (window = 15 frames). Dashed line marks stimulus contact. Related to Fig. 1.

**Fig. S7.**
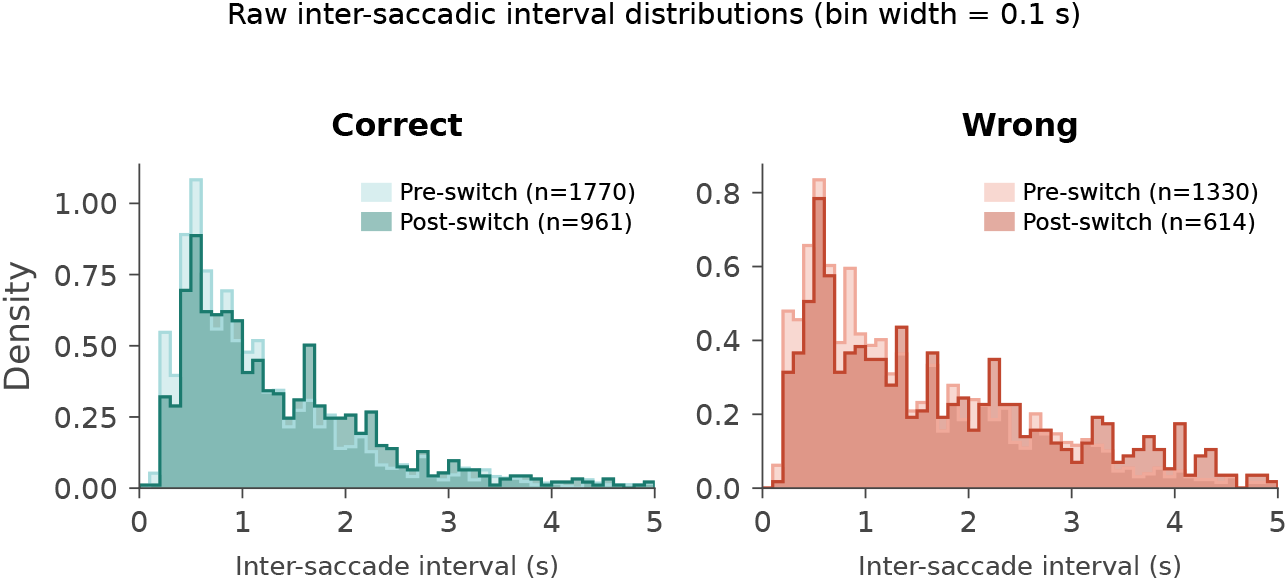
Raw inter-saccadic interval distributions across the rule switch. As in Fig. 5D, but showing the underlying inter-saccadic intervals (ISIs) as unsmoothed histograms (bin width 0.1 s, density-normalised) rather than kernel density estimates, shown separately for correct (left) and wrong (right) trials, pre-switch (light) versus post-switch (dark). The same ISIs as Fig. 5D are plotted (*n* = 1,770 pre / 961 post on correct; 1,330 pre / 614 post on wrong). The plot range is truncated at 5.0 s for visibility; a small number of longer ISIs fall beyond this range (correct: 52 pre / 25 post; wrong: 36 pre / 40 post) and are included in all statistics but not displayed. The post-switch lengthening of ISIs reported in Fig. 5D is visible directly in the unsmoothed distributions. Related to Fig. 5.

